# CAR T-cell and oncolytic virus dynamics and determinants of combination therapy success for glioblastoma

**DOI:** 10.1101/2025.01.23.634499

**Authors:** Martina Conte, Agata Xella, Ryan T. Woodall, Kevin A. Cassady, Sergio Branciamore, Christine E. Brown, Russell C. Rockne

## Abstract

Glioblastoma is a highly aggressive and treatment-resistant primary brain cancer. While chimeric antigen receptor (CAR) T-cell therapy has demonstrated promising results in targeting these tumors, it has not yet been curative. An innovative approach to improve CAR T-cell efficacy is to combine them with other immune modulating therapies. In this study, we investigate *in vitro* combination of IL-13R*α*2 targeted CAR T-cells with an oncolytic virus (OV) and study the complex interplay between tumor cells, CAR T-cells, and OV dynamics with a novel mathematical model. We fit the model to data collected from experiments with each therapy individually and in combination to reveal determinants of therapy synergy and improved efficacy. Our analysis reveals that the virus bursting size is a critical parameter in determining the net tumor infection rate and overall combination treatment efficacy. Moreover, the model predicts that administering the oncolytic virus simultaneously with, or prior to, CAR T-cells could maximize therapeutic efficacy.

## 1 Introduction

Glioblastoma (GBM) is the most aggressive and lethal subtype of primary brain cancer, characterized by a devastating prognosis with a median survival of 12-18 months and limited therapeutic options [55, 74]. The inherent aggressiveness of GBMs is attributed to their rapid proliferation and infiltrative spread into surrounding brain tissue, making them particularly difficult to treat. These features contribute to the tumor’s relentless progression, making it one of the most challenging cancers to manage.

One promising therapeutic strategy that has emerged in recent years is chimeric antigen receptor (CAR) T-cells therapy, where T-cells, taken from the patient or from a donor, are genetically engineered to target antigens expressed on cancer cells [41, 35]. Although CAR T-cells have shown encouraging activity against various solid cancers, including glioblastoma, they are not yet curative. The complexity of the tumor microenvironment, the ability of GBM cells to evade immune responses, and the limitations in CAR T-cell persistence all contribute to the challenges of achieving long-term remission. Because CAR T-cells are a living therapy, they are capable of proliferating, expanding, and adapting over time. These dynamics introduces complex interactions between CAR T-cells and tumor cells, where the kinetics of CAR T-cell expansion and tumor cell killing may vary significantly, influencing the therapeutic outcome. An effective approach to improve CAR T-cell therapy is to combine it with other therapies that may amplify their ability to proliferate, persist, and more effectively target and kill tumor cells. Several such combinations have been studied, including radiation [73, 2], checkpoint-blockade [37, 34], cytokines [79], vaccines [16, 69], among others [3]. The combination of oncolytic viruses with CAR T cells has also been tested in melanoma cells in *in vivo* models, yielding interesting results regarding the interaction between the therapies [30, 31].

In this study, we investigate the combination of 1L13-R*α*2-targeted CAR T-cells with C134 oncolytic virus as a potential therapeutic strategy for GBM. Oncolytic viruses (OVs) are an emerging class of cancer immunotherapies that selectively infect and kill tumor cells, while simultaneously stimulating the host immune system. Currently, a diverse range of OVs has entered different clinical trials for GBM, with promising early results in terms of both safety and efficacy [61, 51]. The unique properties of oncolytic viruses offer several advantages for cancer treatment. For example, in addition to directly inducing lysis of tumor cells, OVs can trigger the release of tumor-associated antigens, which are then presented to the immune system, effectively acting as a vaccine that primes the body’s immune response against the tumor[78]. OVs may enhance CAR T-cell expansion and persistence by creating a more immunologically favorable tumor microenvironment, while CAR T-cells can provide a targeted immune response that augments the anti-tumor effects of the virus.

The complex interactions between cancer cells, CAR T-cells, and oncolytic viruses highlight the need for mathematical modeling to optimize the timing and effectiveness of the combined therapies. Mathematical models are a powerful tool in immunotherapy research [28, 29, 45], enabling the integration of experimental data and providing a quantitative framework for understanding and predicting the dynamics of tumor-immune system interactions. These models also allow for the exploration of different treatment strategies [21, 20, 1]. In recent years, CAR T-cell therapies have drawn significant interest from mathematicians studying a range of tumors [60, 17], including gliomas [65, 6, 66, 47, 7], melanomas [5], and B-cell malignancies [46, 52, 57]. These mathematical models have been used to investigate various cellular mechanisms, such as CAR T-cell activation, cancer cell killing in response to CAR T-cell dose, target antigen expression, and optimal dosing strategies. The dynamic interactions between tumor cells, immune cells, and OVs have also been the focus of many studies [44, 76, 39, 49].

In the context of oncolytic virotherapy, mathematical models have been instrumental in identifying critical parameters [62, 71, 59], generating testable hypotheses, predicting therapeutic outcomes *in silico* [40, 56], and optimizing combination treatments [32, 68, 77]. In both contexts, much of the existing literature has focused on the temporal dynamics of tumor-virus or tumor-virus-immune cell interactions, given the availability of temporal data. Although the combination of CAR T cell therapy and oncolytic virotherapy for solid tumors is widely regarded as promising [58, 53], only a limited number of mathematical models have explored the dynamic effects of this combined therapy. Notably, Mahasa et al. [50] developed a mathematical model incorporating virus-induced synergism to investigate the potential effects of various therapy combinations on tumor cell populations. However, as the authors note, this model serves primarily as a proof-of-concept, and it lacks experimental data to support its conclusions. Additionally, the synergistic effect of the two therapies is directly incorporated into the model, rather than emerging as an emergent property of the system. Thus, while mathematical models have provided valuable insights into the individual and combined effects of CAR T cell therapies and oncolytic virotherapy, there remains a need for more comprehensive models that better capture the complexity of their interactions and provide data-driven predictions to guide clinical applications.

Using an *in vitro* assay, we experimentally evaluated various combinations of CAR T-cell and oncolytic virus treatments against glioblastoma. Using concepts from ecology and epidemiology, we constructed a mathematical model that captures the key interactions between the tumor, CAR T-cells, and oncolytic viruses. This model was then fitted to our experimental data to provide a quantitative framework for understanding the dynamics of these therapies. By using this model, we were able to explore how the timing and coordination of CAR T-cell and OV treatments impact their combined therapeutic efficacy. Our analysis revealed highly nonlinear relationships between the treatment schedules and the outcomes, demonstrating that small changes in the timing or sequence of the therapies can significantly alter their effectiveness. This modeling approach provides deeper insights into the complex dynamics of tumor-immune interactions, offering potential strategies to optimize the combination of CAR T-cell and OV therapies for glioblastoma. By better understanding how these therapies interact, our approach provides valuable strategies for improving therapeutic outcomes and guiding clinical decisions.

This manuscript is organized as follows: first we describe the experimental methods and data, followed by a detailed motivation and description of the mathematical model that includes both CAR T-cells and OV. Then we examine sub-models which include only CAR T-cells or OV individually. We fit the sub-models to respective monotherapy experimental data, and then use those parameters to fit the full model to the combination therapy data. We analyze and interpret the parameter values as they vary by treatment dose, and then to explain the benefit of combination therapy. Finally, we use the fully parameterized model to study the effect of therapy timing and to predict dynamics of different treatment combinations.

## 2 Methods

### 2.1 Experimental design

#### 2.1.1 Cell lines

Low-passage primary brain tumor (PBT) lines were derived from GBM patient nr.30 (PBT030) that had undergone tumor resections at City of Hope as previously described [8, 9, 10]. PBT030 cells were maintained in Neurosphere conditions [11] and passaged 2 times a week. Cells were kept low passage and used after 1 week but before 2 months from thawing. PBT030 cells have endogenously high IL-13R*α*2 expression and are naturally targeted by the CAR T-cells used.

Chimeric antigen receptor T-cells targeting IL-13R*α*2 were obtained from healthy donor T-naïve/T-memory cells (CD25 and CD14 depleted, CD62L enriched Peripheral Blood Cells) activated with CD3-CD28 Dynabeads™(Invitrogen), transduced with IL-13(EQ)BB*ζ*/CD19t lentivirus, and expanded in X-Vivo media with 50 U/mL rhIL-2 and 10 ng/mL rhIL-15 for 14 to 18 days [8]. Expression of lentiviral construct was checked by flow cytometry.

#### 2.1.2 Oncolytic Virus

Research grade herpes simplex virus (HSV)-based oncolytic virus C134 was obtained from the laboratory of Dr. Cassady at Nationwide Children’s Hospital in Columbus, OH [14, 15, 33]. Aliquoted and titered vials of the virus in PBS with 10% glycerol, that passed bacterial and fungal testing, were shipped and received frozen and kept at -80°C. Virus aliquots were thawed as needed and kept on ice for the time necessary for the experimental use.

#### 2.1.3 Infection assay

Single cell suspensions of primary brain tumor (PBT) lines in neurosphere media (50:50 mix of DMEM and F12 medias with 1% glutaMAX supplement, 2% B-27 supplement, 1.5% HEPES buffer, and 1,000 USP/mL Heparin) with 10% FBS were plated at 1 ×10^5^ cells per 100 *µ*L/well in a flat bottom 96 well plate overnight to allow for attachment to the plate. The next day, OV stock in 10% glycerol was diluted in neurosphere media with 10% FBS to achieve the necessary number of plaque forming units (PFU) per 20 *µ*L for a multiplicity of infection (MOI, i.e., PFU/cell number) of either 1.0, 0.1, 0.01, 0.001, or 0.0005. Culture media (60 *µ*L/well) was removed prior to the addition (20 *µ*L/well) of the diluted OV, or controls of glycerol alone or media alone. Each condition was assayed in duplicate wells. The plate was incubated at 37°C, 5% CO_2_ for 2 hours, and subsequently washed 3 times with media, leaving 150 *µ*L of media prior to returning it to the incubator. A 48 hours post infection, cells were harvested mechanically and by using Accutase, and processed for flow cytometric analysis. In brief, cells were stained with Fixable Viability Dye eFluor™506 and anti-HSV-gD antibody in FACS stain solution (FSS, HBSS with 1 g/L Sodium azide and 2% FBS). Samples were then run and analyzed on a MACSQuant10 (Miltenyi) instrument, and percentages of viable infected (i.e., HSV-gD+ and GFP+) cells were calculated via FlowJo software v10.8.0 or later (FlowJo, LLC, Ashland, OR).

#### 2.1.4 Killing Assay

To evaluate the killing kinetics of the OV C134 and IL-13R*α*2-CAR T-cells against PBT030, we used the xCELLigence analyzer (ACEA Bioscience) [63] (Figure 1–A). The xCELLigence system uses electrical impedance to non-invasively measure adherent cell density of a specialized 96 well E-Plates. This measurement is translated in a dimensionless number referred to as Cell Index (CI) that is strongly positively correlated with the number of cells in the well and, thus, is used as a measure of cell number [75]. To better reveal any potential benefit of the combination treatment we used suboptimal doses of both OV C134 and IL-13R*α*2-CAR T-cells.

**Figure 1:**
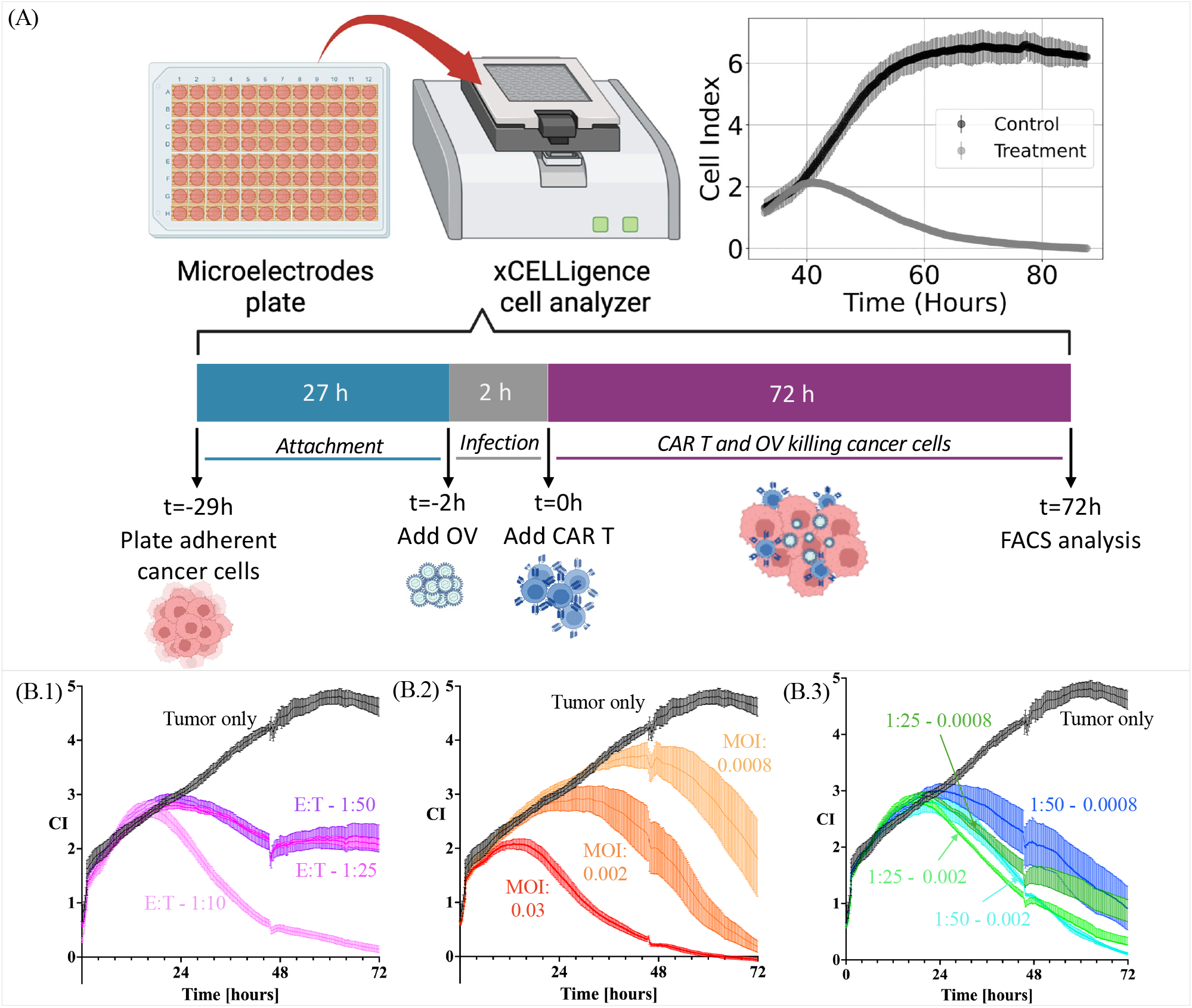
Experimental setting and data collection. (A) Diagram of experimental procedure using microelectrode plates in an xCELLigence cell analyzer system, adapted from [12]. (B) Experimental data collected from glioblastoma tumor cells treated with (B.1) CAR T-cells only, (B.2) oncolytic virus only, or (B.3) CAR T-cells and oncolytic virus simultaneously for different effector to target ratios and initial virus concentrations measured with the xCelligence platform in units cell index (CI).

Neurosphere media with 10% FBS was used for all xCELLigence experiments, and it will be referred as *media* for this session. On the first day (t=-29h), plates were blanked with 50 *µ*L of media alone on the machine to establish baseline reading. PBT030 neurospheres were mechanically and enzymatically (Accutase) removed from the flask and dissociated, final cell straining was done to ensure single-cell status. Cells were counted, and resuspended at a concentration 2 ×10^4^ cells per well with a final volume 140 *µ*L per well. We had 3 replicate wells per condition, and no outside wells in the plates were used to prevent edge effect biases, but they were filled with media to help prevent evaporation in experimental wells (moat). Cells were left to attach for 1h at RT outside the xCELLigence machine and for about 23h additional hours at 37°C inside the machine. CI was set to record every 15 minutes for the rest of the experiment.

After initial incubation, the plate analysis was paused when impedance reached a cell index of ≈0.9-1, and 90 *µ*L of media was removed prior to the addition (20 *µ*L/well) of OV C134 that had been diluted in media to achieve a final MOI (PFU/cell number) of either 0.03, 0.002 or 0.008, or glycerol alone (at the same concentration as the MOI of 0.03 condition) as a control. The initial cell number of 2 ×10^4^ was used for the calculation as we assumed minimal o no growth during the attachment phase. The plate was placed back on the xCELLigence instrument within a 37°C incubator and impedance measurements collected every 15 minutes were resumed for 2 hours. Plate analysis was again paused, the plates were washed by adding and removing ≈75% of media volume twice, and then IL-13R*α*2 CAR T-cells in neurosphere media with 10% FBS were added at 1:10, 1:25 or 1:50 CAR+ effector to target (E:T) cell ratios (based on initial number of plated tumor cells). The plate was placed back on the xCELLigence instrument within a 37°C incubator (t=0 h) and recording of impedance every 15 minutes was resumed. Around 2 days post treatment E-plates were removed, and the moat wells were re-filled to prevent evaporation in the experimental wells. Three days after treatment, the recordings were stopped and all electrical impedance data were collected from the xCELLigence instrument. Analysis and plotting of the raw data was done using Excel and Prism, as shown in Figure 1–B. For this study, the tumor *area under the curve* (AUC) was quantified for both data and model simulations as a quantity of interest to both capture the accuracy of the fits and provide a common metric for comparing different treatment scenarios [67].

### 2.2 Mathematical model

To provide new insights into the biological data described in Section 2.1, we develop a mathematical model to analyze the interactions between glioblastoma tumor cells and the two therapeutic agents: CAR T-cells and oncolytic viral particles. This model draws inspiration from variations of classical ecological and epidemiological approaches to assess the effects of monotherapy treatments on tumor cells. Specifically, we utilize susceptible-infected-recovered (SIR) type models to describe the spread of infection [70] and Lotka–Volterra equations to model interactions between the tumor cells, CAR T-cells, and virus [27, 54].

Our model includes four populations that are assumed to be well mixed: (non-infected) tumor cells (*T*), infected tumor cells (*I*), oncolytic virus (*V*), and CAR T-cells (*C*), whose interactions are illustrated in Figure 2. Precisely, in the absence of treatment, tumor cell growth is limited only by space and nutrient (culture media) in the *in vitro* culture system and, thus growing logistically. When oncolytic virus particles are introduced into the environment, they infect tumor cells, converting them to the infected population. The infected tumor cells lose their ability to proliferate and eventually undergo lysis, releasing new viral particles into the environment. The viral burst phenomenon increases the viral particles in the system and counteracts natural clearance of virus. Conversely, the administration of CAR T-cells leads to interactions with tumor cells that can either stimulate or exhaust CAR T-cells, which are also subject to natural apoptosis. When both therapies are applied simultaneously, complex dynamics can emerge, involving both competition and cooperation between CAR T-cells and the oncovirus. Notably, CAR T-cells can eliminate both tumor and infected tumor cells, while they themselves can become infected by the virus, compromising their cytotoxic capabilities.

**Figure 2:**
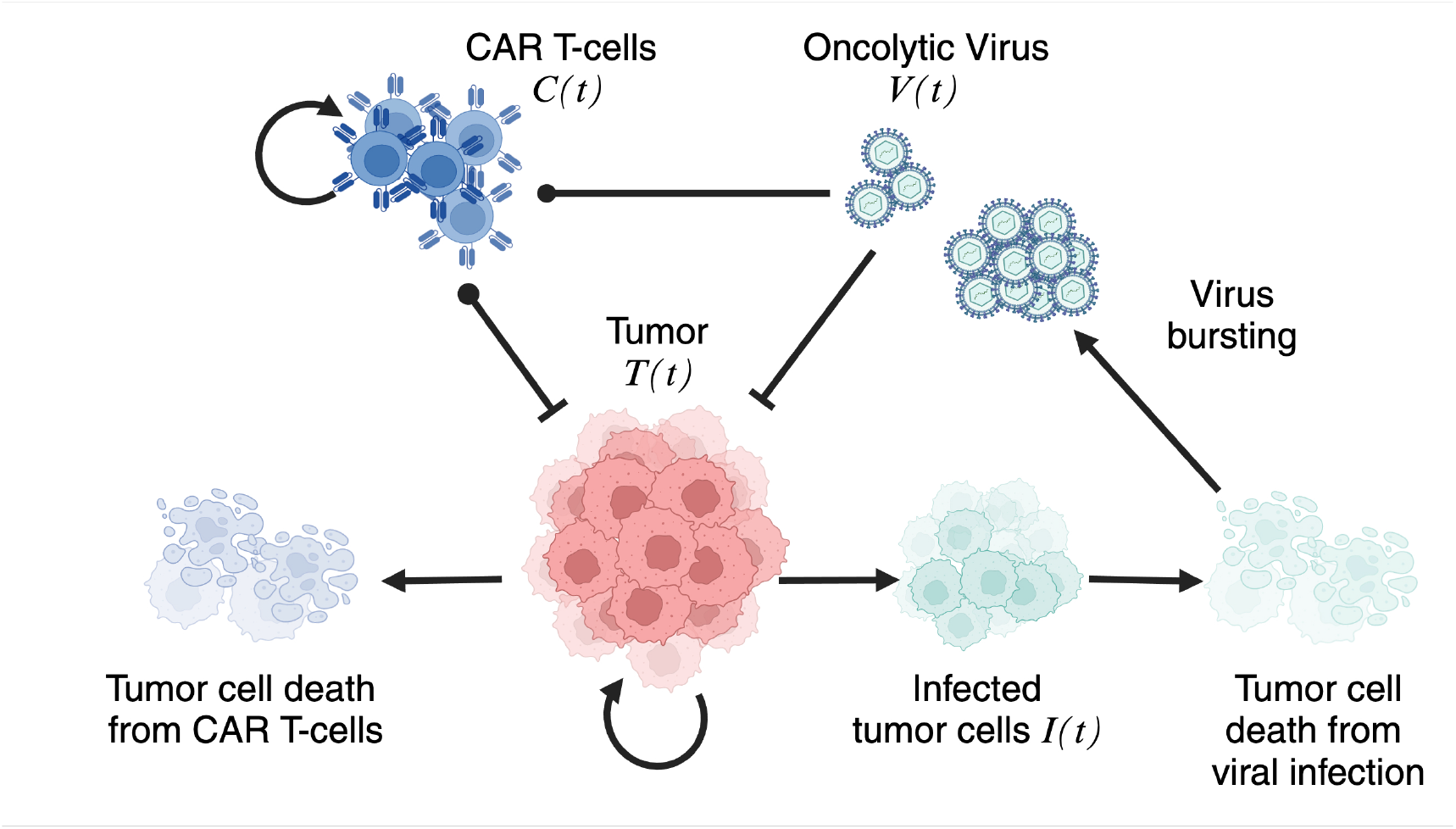
Schematic of the model. A schematic representation of the interactions between the different populations in the model is provided. Loop arrows represent cell proliferation, while normal arrows indicate the conversion of one population into another. Vertical bars and black dots signify negative effects, such as cell killing or exhaustion, exerted by one population on another.

The interactions outlined can be formalized through the following system of ordinary differential equations (ODEs):

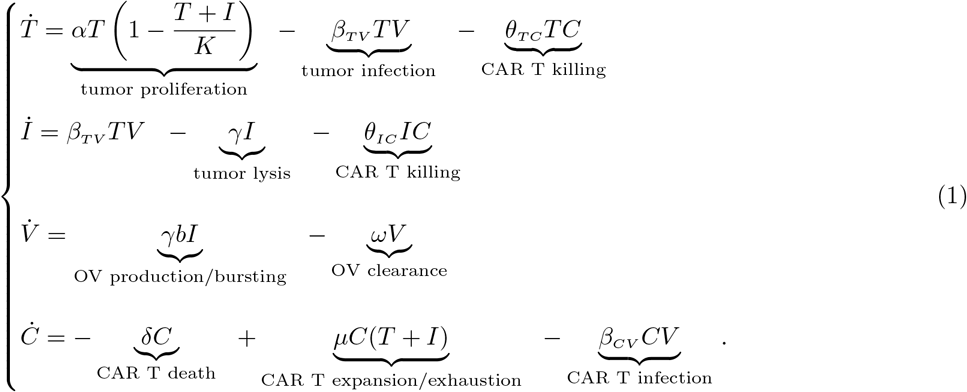

This system models the dynamics between tumor cells, infected cells, oncolytic virus, and CAR T-cells where *α* is the growth rate of tumor cells, and *K* is the tumor carrying capacity. *β* is the infection rate of either tumor cells (*β*_*TV*_) or CAR T-cells (*β*_*TC*_) due to the virus activity, and *θ* represents the CAR T killing rate of either tumor (*θ*_*TC*_) or infected tumor cells (*θ*_*IC*_). The remaining parameters refer to the lysis rate of infected tumor cells (*γ*) or CAR T-cells (*δ*), the virus clearance rate (*ω*), and the expansion/exhaustion rate of CAR T-cells due to the interactions with the tumor cells (*µ*).

The burst size of the virus is given by *b* which is the number of new viral particles released into the environment upon lysis of an infected tumor cell. The burst size parameter is crucial for accurately modeling viral dynamics and its influence on CAR T-cell and tumor behavior. A higher burst size *b* results in the release of more viral particles with each lysis event. We summarize the model parameters with their meaning and their units of measure in Table 1.

**Table 1:** Model parameter. Summary of the model parameters with their description and their unit of measure. *h* : hours, CI: cell index, MOI: virus multiplicity of infection.

| Parameter | Description | Unit |
| --- | --- | --- |
| $\alpha$ | tumor proliferation rate | $h^{-1}$ |
| $K$ | tumor carrying capacity | CI |
| $\beta_{TV}$ | tumor infection rate | $(\text{MOI} \cdot h)^{-1}$ |
| $\beta_{CV}$ | CAR T infection rate | $(\text{MOI} \cdot h)^{-1}$ |
| $\theta_{TC}$ | CAR T killing rate of tumor cells | $(\text{CI} \cdot h)^{-1}$ |
| $\theta_{IC}$ | CAR T killing rate of infected tumor cells | $(\text{CI} \cdot h)^{-1}$ |
| $\gamma$ | infected tumor cell lysis rate | $h^{-1}$ |
| $b$ | virus burst size | $\text{MOI} \cdot (\text{CI})^{-1}$ |
| $\omega$ | virus clearance rate | $h^{-1}$ |
| $\delta$ | CAR T-cell death rate | $h^{-1}$ |
| $\mu$ | expansion/exhaustion CAR T-cell rate | $(\text{CI} \cdot h)^{-1}$ |

The experiments performed examine both the monotherapy scenarios (using either CAR T-cells, or oncolytic virus alone) and the combined approach. We first analyze two simplified versions of model (1), before presenting the results for the full model.

We define the **CAR T-cell module** as the following reduced system of ordinary differential equations:

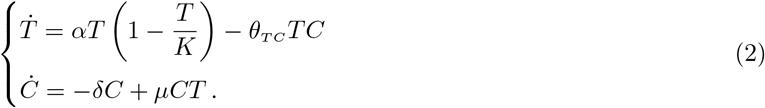

Originally introduced in [65] as the CARRGO model, this system employs classical predator–prey dynamics to investigate the kinetics of CAR T-cell killing in gliomas. In this model, we assume that *α, K, θ*_*TC*_, and *δ* are positive parameters. The parameter *µ* can take either positive or negative values, allowing us to capture both the stimulation and exhaustion of CAR T-cells as they interact with tumor cells. This flexibility provides insight into the varying effects of CAR T-cell engagement in the therapeutic context.

For the oncolytic virus therapy alone, we define the **OV module** as the following reduced system of ordinary differential equations:

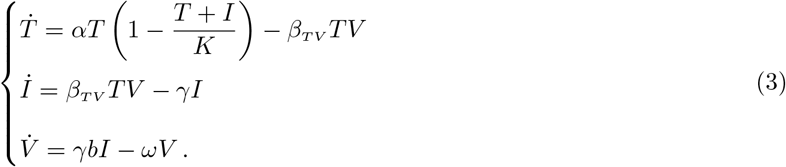

This model, inspired by classical susceptible-infected-recovered dynamics, characterizes the infection process of tumor cells resulting from the introduction of viral particles into the system. In this framework, all model parameters are assumed to be positive. A qualitative analysis of both CAR T-cells and OV modules is provided in the Supplementary Material S.1. Numerical simulations of the mathematical models were performed using a self-implemented code in Matlab.

### 2.3 Model fitting to experimental data

In all experiments, the initial 24 hours of the time-series capture the process of cell attachment to the bottom of the plate. Following the approach outlined in [65], we omit this attachment phase from our analysis, as we are primarily interested in cell growth and interaction kinetics. We utilize data from 24 to 96 hours post seeding to estimate and analyze the model parameters. The fitting procedure used to estimate model parameters uses a self–implemented code in MATLAB, based on the Particle Swarm Optimization (PSO) algorithm for global optimization [25, 42]. PSO is a stochastic global optimization procedure inspired by biological swarming and recently used for parameter estimation in a variety of initial value problems across cancer research and systems biology [22, 64, 38, 13]. These optimization procedures are used to minimize the sum–of–squares error (SSE) between measured and predicted tumor cell populations. Specifically, the carrying capacity *K* is estimated by fitting a logistic growth model to three replicates of time–series data of untreated cancer cells. Next, we estimate the parameters of the **CAR T-cell module** (*θ*_*TC*_, *δ, µ*) by fitting model (2) to the CAR T-cell monotherapy experiment dataset with tumor cells treated with CAR T-cells at different effector to target ratios of 1:10, 1:25, and 1:50. For fitting the parameters of the **OV module** (*β*_*TV*_, *γ*) we use the OV monotherapy experiment dataset of tumor cells treated with oncolytic virus at 0.0008, 0.002, 0.03 MOI initial concentrations. Finally, we use the CAR T-cell plus OV combination therapy experiment dataset of tumor cells treated with CAR T-cells and oncolytic virus to fit the remaining parameters of model (1) (*θ*_*IC*_, *β*_*IC*_), along with *β*_*TV*_ and *θ*_*TC*_, to investigate the potential cooperative or competitive effects of the combined therapies. All experimental conditions were repeated in triplicate. Model parameters were fit to each of the replicates and the mean value and standard deviation of the obtained estimations were considered.

## 3 Results

From the experimental results of this study, we observe that high concentrations of either CAR T-cells or the oncolytic virus alone can effectively eradicate the tumor population *in vitro*. However, their efficacy diminishes at lower therapy doses. In contrast, combinations of the two treatments, even at lower concentrations, yield better outcomes than monotherapy alone.

We utilize the mathematical model to provide insights into these biological findings and to serve as a predictive *in silico* tool for testing drug combination efficacy in a variety of different scenarios, including low and experimentally challenging drug concentrations and different timing of therapy administration. This approach not only enhances our understanding of the interactions between the therapies but also aids in optimizing treatment strategies for improved therapeutic outcomes.

### 3.1 Nonlinear CAR T dose–dependent dynamics

We first investigate the CAR T monotherapy case, described by model (2), analyzing the impact of varying the CAR T-cell dose, quantified by the effector to target ratio. The tumor response is assessed for CAR T-cells at E:T ratios of 1:10, 1:25, and 1:50, as illustrated in Figure 1–B.1. The results of data-fitting are presented in Figure 3, where panels A, B, and C show the temporal evolution of model (2) (tumor cells in red, CAR T-cells in blue) compared to the original data (mean ± std, in black) for the three different E:T ratios. Panel D compares the AUC for all datasets and experimental conditions.

**Figure 3:**
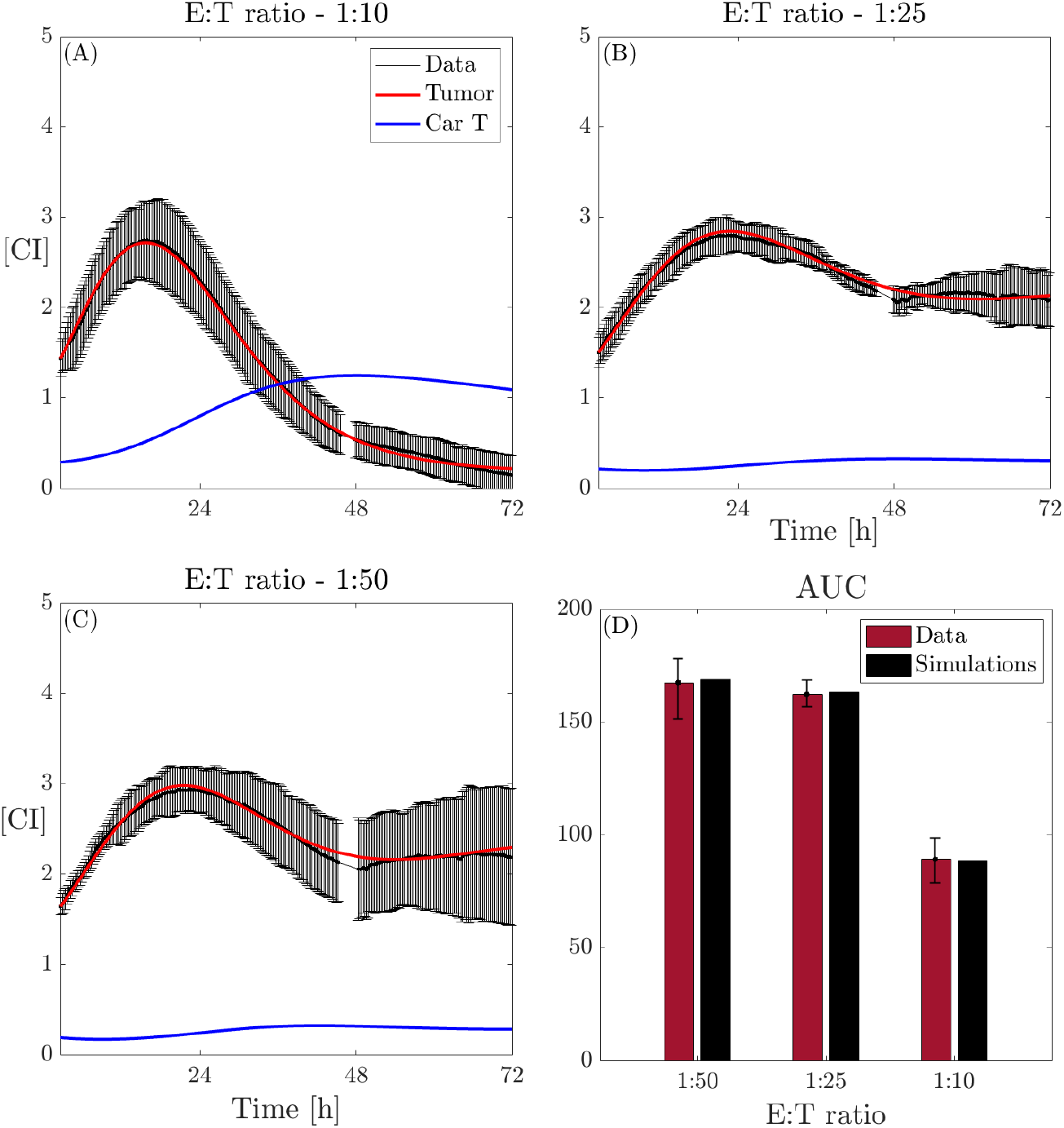
Dynamics of model (2) and *in vitro* CAR T-cell and glioma cell data. Dynamics of cancer cells along with the model fits (black: data, red line: tumor cells, blue line: CAR T-cells) with effector to target ratios (A) 1:10, (B) 1:25, and (C) 1:50. (D) Comparison of AUC obtained from the *in vitro* (red) data and *in silico* (black) results. Mean *±* std is obtained from the three replicates of the same experiment.

Upon analyzing the parameter values, we find that the CAR T expansion/exhaustion rate *µ* is consistently small and positive across all CAR T-cell doses, decreasing as the E:T ratio increases. A similar trend is observed for the CAR T-cell death rate *δ*, which is higher than *µ* in at lower CAR T-cell doses. This finding appears to contrast with the results presented in [65]. However, this phenomenon can be explained by noting that the constant *µ* reflects two biological processes: expansion and exhaustion, thus representing the net effect of these opposing dynamics. In high E:T ratios, the injected CAR T-cells undergo both enhanced expansion and increased exhaustion, leading to a delicate balance that results in low *µ* and *δ* values. Biologically, higher E:T ratios lead to faster dynamics of activation and exhaustion of CAR T-cells, contributing to a more rapid response. Nevertheless, the changes observed in *µ* manifest inversely in *δ*, resulting in a higher net CAR T-cell expansion rate over time at elevated E:T ratios, aligning with the findings of [65]. The killing rate parameter *θ*_*TC*_ exhibits a nonlinear relationship with the CAR T-cell dose. At higher doses, *θ*_*TC*_ shows a negative correlation with the dose, while at lower doses, the correlation becomes positive, indicating an optimal CAR T-cell killing capability at intermediate doses. The results of this analysis are summarized in Figure 4, where panels A, B, and C show variation of *θ*_*TC*_, *µ*, and *δ*, respectively, across the different E:T ratios. Average values for the estimated parameters across different doses are provided in the Supplementary Table S.1 in the Supplementary Material S.2.

**Figure 4:**
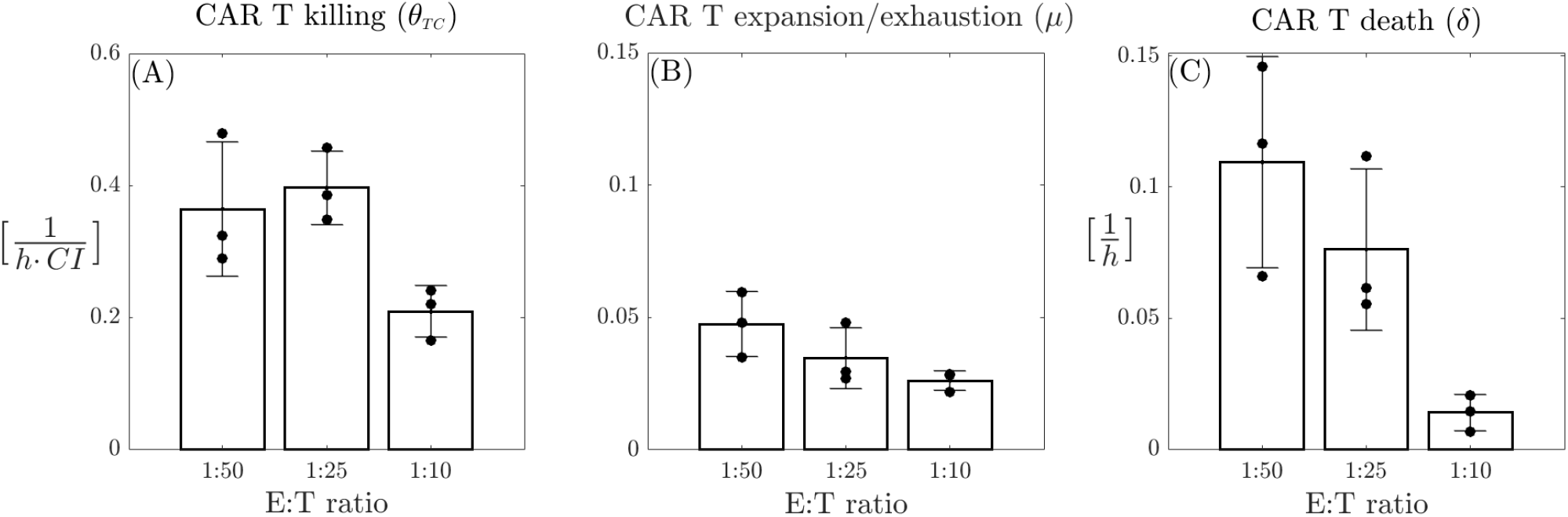
Comparisons of model (2) parameter with CAR T-cell dose. (A) CAR T killing rate (*θ*_*TC*_); (B) CAR T expansion/exhaustion rate (*µ*); (C) CAR T death rate (*δ*). Each parameter is plotted for the three different effector to target ratios (1:10, 1:25, and 1:50). Bars represent the mean values of the parameters, while error bars indicate the standard deviation. Black bullets denote the individual values for each of the three replicates from the experiments.

### 3.2 Impact of viral dynamic parameters on therapy efficacy

We investigate the oncolytic virus monotherapy case, described by model (3), by varying the initial concentration of the virus to analyze the impact of viral dynamic parameters, namely, burst size *b*, infection rate *β*_*TV*_, and clearance rate *ω* on therapy outcomes. Tumor response is evaluated for initial OV concentrations of 0.03 MOI, 0.002 MOI, and 0.0008 MOI, as illustrated in Figure 1–B.2.

#### 3.2.1 Virus burst size analysis

The burst size *b* is a crucial parameter that quantifies the total number of virions produced by an infected cell. While several efforts have been made to establish reliable procedures for identifying this parameter across different viruses [18, 24, 26], it remains somewhat ambiguous. Typically, the range of variation for burst size is estimated to be between [0.1 ™10^4^] PFU/cells [19]. To understand how variability in the burst size *b* may impact the dynamics of the system under investigation, we begin by analyzing whether and how the estimates of tumor related parameters in model (3), namely, *α, β*_*TV*_, *γ*, are influenced by changes in *b*. For this analysis, we first convert the range of variability of *b* from PFU/cells to MOI/CI. Considering the relation between cell number and CI provided in [65] and the relation 1MOI = 20 *·* 10^3^ PFU, it is possible to express *b* in unit MOI/CI recovering the following range of variability of the bursting size *b* ∈ [0.025, 2500] MOI/CI.

In our analysis of the three different scenarios, i.e., initial virus dose of *V*_0_ = 0.03, 0.002, 0.0008 MOI, we find that the only parameter consistently affected by variations in the burst size *b* is the tumor infection rate *β*_*TV*_. No significant changes are observed in the estimates for *α* and *γ* (see Supplementary Figure S.3 and Supplementary Table S.4 in the Supplementary Material S.3). As shown in Figure 5-A, the estimation of *β*_*TV*_ varies dramatically across several orders of magnitude in response to changes in *b* within the range of [0.025, 2500]. The higher the burst size value, the smaller the infection tumor rate. Furthermore, it is possible to analytically demonstrate that *β*_*TV*_ and *b* are logarithmically related, a trend that is also evident in Figure 5–B. Specifically, given *ω*, and considering the known quantities 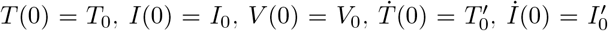, and 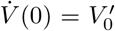, we can evaluate model (3) at time *t* = 0 as

**Figure 5:**
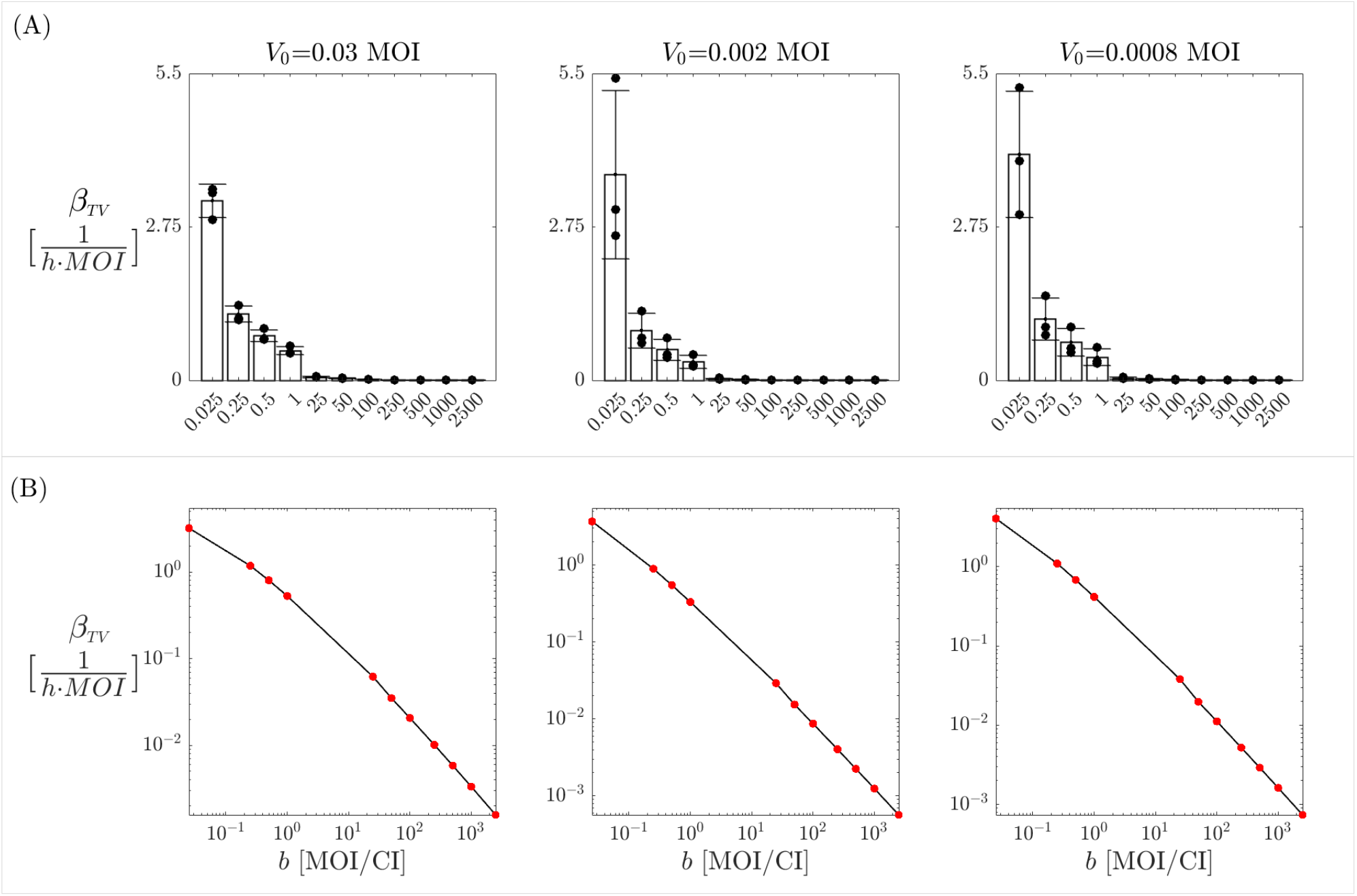
Burst size effects on tumor infection rate. (A): Estimation of the parameter *β*_*TV*_ for various burst size values *b* within the range of [0.025, 2500]. (B): a log–log plot illustrating the inverse relationship between *β*_*TV*_ and *b*. Red dots represent the specific pairs of *b, β*_*TV*_ obtained from the analysis, while the black curve indicates their linear interpolation. In both rows, the columns correspond to the three different virus doses.

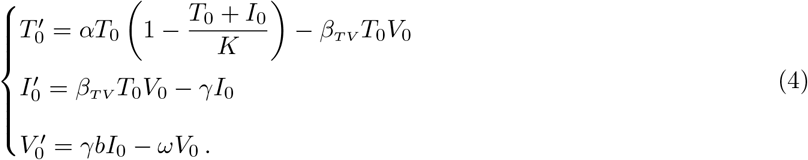

From the second and third equations in (4), we get

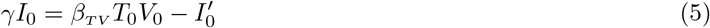

and

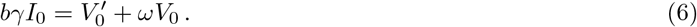

Substituting (5) into (6) we obtain

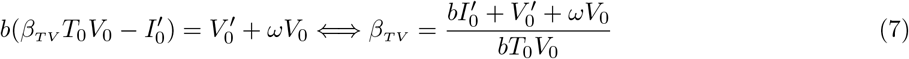

which is well defined since *T*_0_, *V*_0_, *b* ≠0. Assuming that *I*^*′*^(0) = 0 and applying the logarithm to both side of equation (7) we have

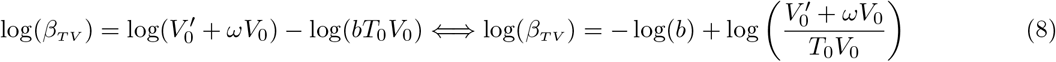

Relation (8) provides an analytical expression for the logarithmic relationship between *b* and *β*_*TV*_, shown in Figure 5–B. Thus, these results underscore the significant impact that the virus–specific burst size can have on the model dynamics, emphasizing the importance of obtaining accurate estimates from experimental data. Since the values of the proliferation rate *α* and the apoptosis rate *γ* remain constant despite variations in the burst size (see Supplementary Figure S.3 and Supplementary Table S.4 the Supplementary Material S.3), we calculate their average across the different burst size values, as presented in Supplementary Table S.2 the Supplementary Material S.2.

By fixing these parameter values, we conduct a second fitting step for the tumor infection rate *β*_*TV*_, varying the burst size *b* within the interval [0.025, 2500]. We then average the results across the different *b* values. The resulting average estimations for *β*_*TV*_ are illustrated in Figure 6. The results in Figure 6 reveal that the infection rate *β*_*TV*_ exhibits a nonlinear relationship with the virus dose. Specifically, it is lower at low virus doses, increases with *V*_0_, and then exhibits a saturation effect as the doses increase. This behavior can be attributed to the saturation of virus particles in the environment, which becomes fully occupied, limiting any further increase in infective capability. The results of the data fitting, obtained using these estimates for the parameters involved in model (3), are presented in Figure 7, where panels A, B, and C show the temporal evolution of model (3) compared to the original data for the three different initial virus doses, while panel D compares the AUC for all datasets and experimental conditions.

**Figure 6:**
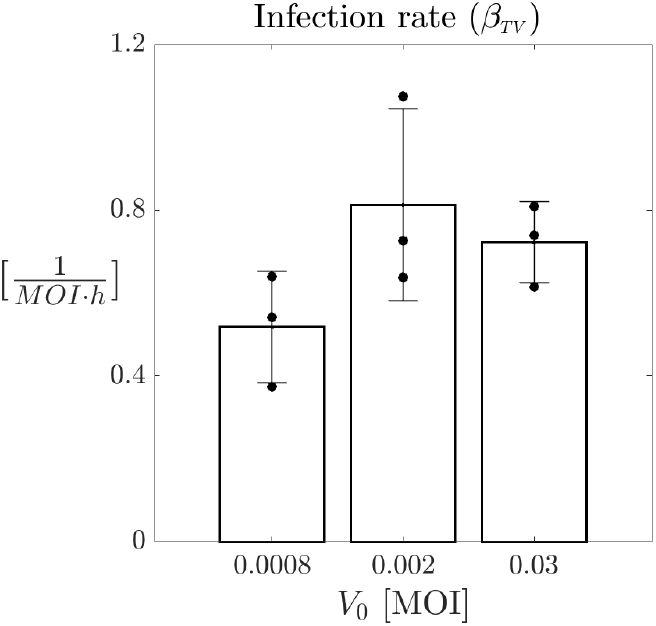
Comparisons of model (3) parameter *β*_*TV*_ with virus dose. Estimations of the tumor infection rate are plotted for the three different initial virus concentration 0.03, 0.002, 0.0008 MOI. Each estimation is obtained as an average across the different *b* values in the range [0.025, 2500]. Bars represent the mean values of the parameters, while error bars indicate the standard deviation. Black bullets denote the individual values for each of the three replicates from the experiments.

**Figure 7:**
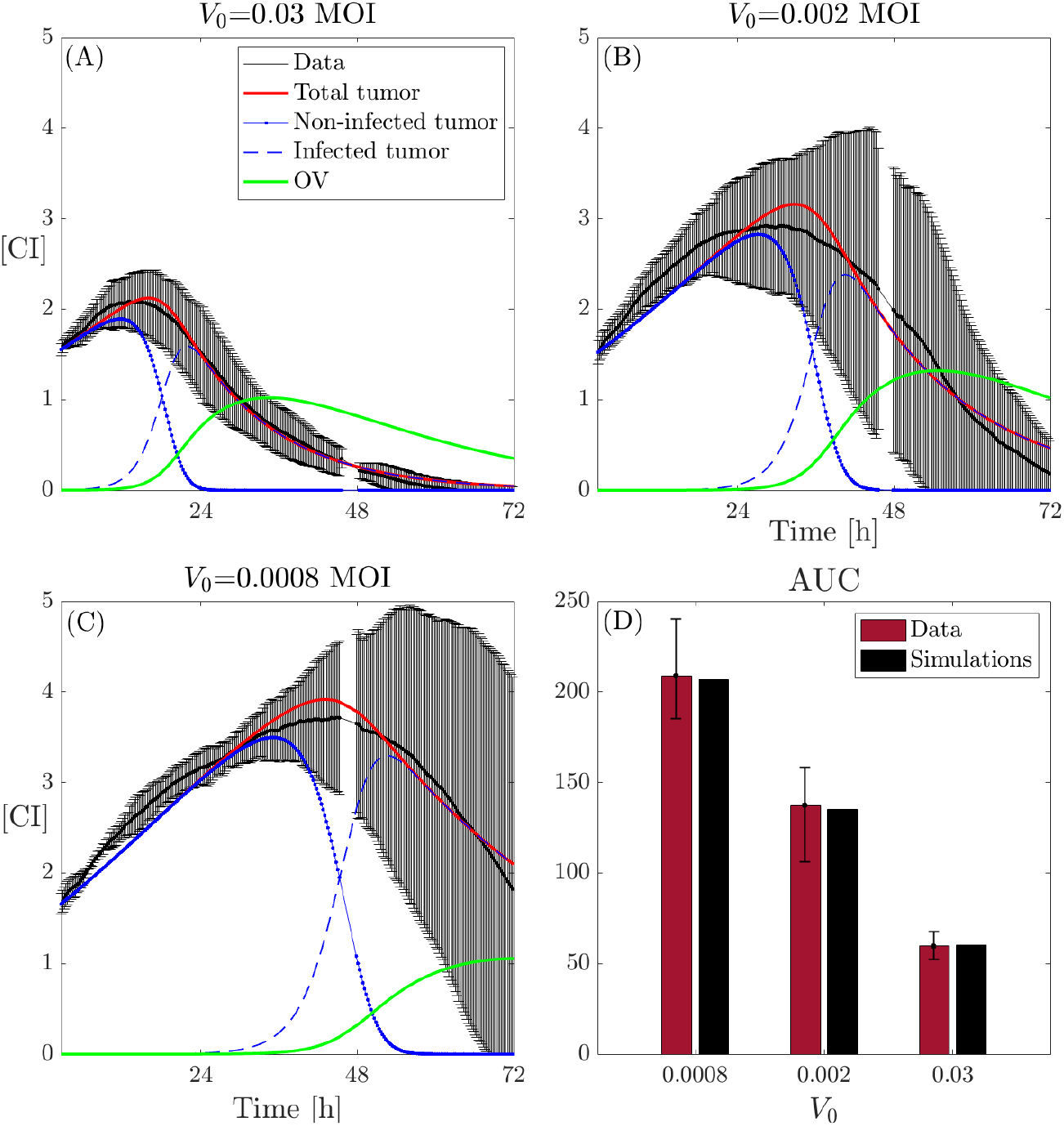
Dynamics of model (3) and *in vitro* OV and glioma cell data. Dynamics of cancer cells along with the model fits (black: data, red line: total tumor cells, blue dashed–dot line: non-infected tumor cells, blue dashed line: infected tumor cells, green line: OV) with initial virus dose (A) 0.03 MOI, (B) 0.002 MOI, and (C) 0.0008 MOI. (D) Comparison of AUC obtained from the *in vitro* (red) data and *in silico* (black) results. In all the cases the burst size is *b* = 25 MOI/CI and the clearance rate is *ω* = 0.05 *h*^−1^.

#### 3.2.2 Infected cell fraction as a function of OV parameters

We analyze the *in vitro* experimental data presented in the Supplementary Figure S.4–A (in black) (see Supplementary Material S.3) related to the fraction of infected tumor cells observed in culture 24 and 48 hours after virus administration. Specifically, the fraction of infected cells *I*(*t*) relative to the total tumor cell population *T* (*t*) + *I*(*t*) is evaluated for two different replicates of the same experiment at both time points 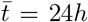and 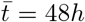, across five different virus doses: *V*_0_ = 0.0005, 0.001, 0.01, 0.01, 1 MOI. Using a cubic interpolation, we estimate the mean fraction of infected cells for *V*_0_ = 0.0008, 0.002, 0.03 MOI and calculate the corresponding standard deviation, which is depicted in red in Supplementary Figure S.4–A. Subsequently, we utilize model (3) to conduct several *in silico* experiments, assessing the effects of the virus–specific parameters, burst size (*b*) and clearance rate (*ω*), on the fraction of infected tumor cells. We then compare these results with the interpolated data, as shown in Supplementary Figure S.4–B.

Our results highlight that, in all scenarios analyzed, variations in at least one of the two virus-specific parameters lead to significant fluctuations in the estimated fraction of infected cells. Notably, for high initial virus concentrations, such as *V*_0_ = 0.03 MOI, both parameters exert a substantial influence on model outcomes, particularly over extended time periods. This underscores the importance of obtaining precise estimates of these parameters from experiments to enable reliable predictions of potential outcomes. Despite the considerable variability in model results, our model effectively replicates the experimental observations across all scenarios. As shown in Supplementary Figure S.4–B, the mean and standard deviations of the experimental data (represented by red solid and dashed lines, respectively) remain within biologically plausible ranges for both *b* and *ω*.

### 3.3 Effects of combined therapy protocols

Finally, we perform a comprehensive investigation of the combined therapy scenario as outlined in model (1). This study aims to elucidate the potential effects of cooperation or competition between the two therapeutic treatments, shedding light on how these interactions can influence treatment outcomes. By varying the initial conditions of the model—such as the time of administration and the concentrations of the treatments—we seek to generate predictions that could inform future experiments, and pre-clinical and clinical strategies. This analysis is based on a dataset comprising four sets of experiments, each with three replicates. This dataset is designed to evaluate the tumor response to treatment across four specific combinations of initial OV concentrations and E:T ratios: E:T=1:50 and *V*_0_ =0.0008, E:T=1:50 and *V*_0_ =0.002, E:T=1:25 and *V*_0_ =0.0008, and E:T=1:25 and *V*_0_ =0.002. The dataset is illustrated in Figure 1–B.3. By analyzing these combinations, we aim to identify patterns and draw meaningful conclusions about the interplay between the two therapies, ultimately enhancing our understanding of how to optimize treatment protocols for better patient outcomes.

#### 3.3.1 Combination therapy synergy

Based on the dataset described, we conducted a data fitting process, the results of which are illustrated in Figure 8. Figure 8-A presents the temporal evolution of model (1) alongside the original data (mean ±std, in black) for four different combinations of initial OV and E:T ratios (A.1-4), as well as the comparison of the AUC values obtained from the experiments (A.5, red) and the numerical simulations (A.5, black). Figure 8-B presents the estimated parameters of model (1) across the four therapeutic combinations. Analysis reveals that the CAR T-cell killing rates for both non-infected tumor cells (*θ*_*TC*_) and infected tumor cells (*θ*_*IC*_) consistently increase with higher initial virus concentrations and E:T ratios. Notably, CAR T-cells demonstrate a significantly greater capacity to kill infected tumor cells compared to non-infected tumor cells. This indicates a synergistic effect between the two therapies, as viral particles sensitize tumor cells to death by CAR T-mediated killing [36]. Moreover, the infection rate *β*_*TV*_ shows a tendency to assume larger values for higher E:T ratios. This may suggest that the combination of treatments enhances the infection effects as well. Interestingly, we noted a slight decrease in *β*_*TV*_ between *V*_0_ = 0.0008 and *V*_0_ = 0.002, indicating that the point of infection saturation (relative to the initial *V*_0_ concentration) is reached at lower doses when both therapies are combined with respect to the monotherapy results (shown in Figure 6). These findings are summarized in Figure 8-B, while the average values for all the estimated parameters across different doses presented in the Supplementary Table S.3 in Supplementary Material S.2.

**Figure 8:**
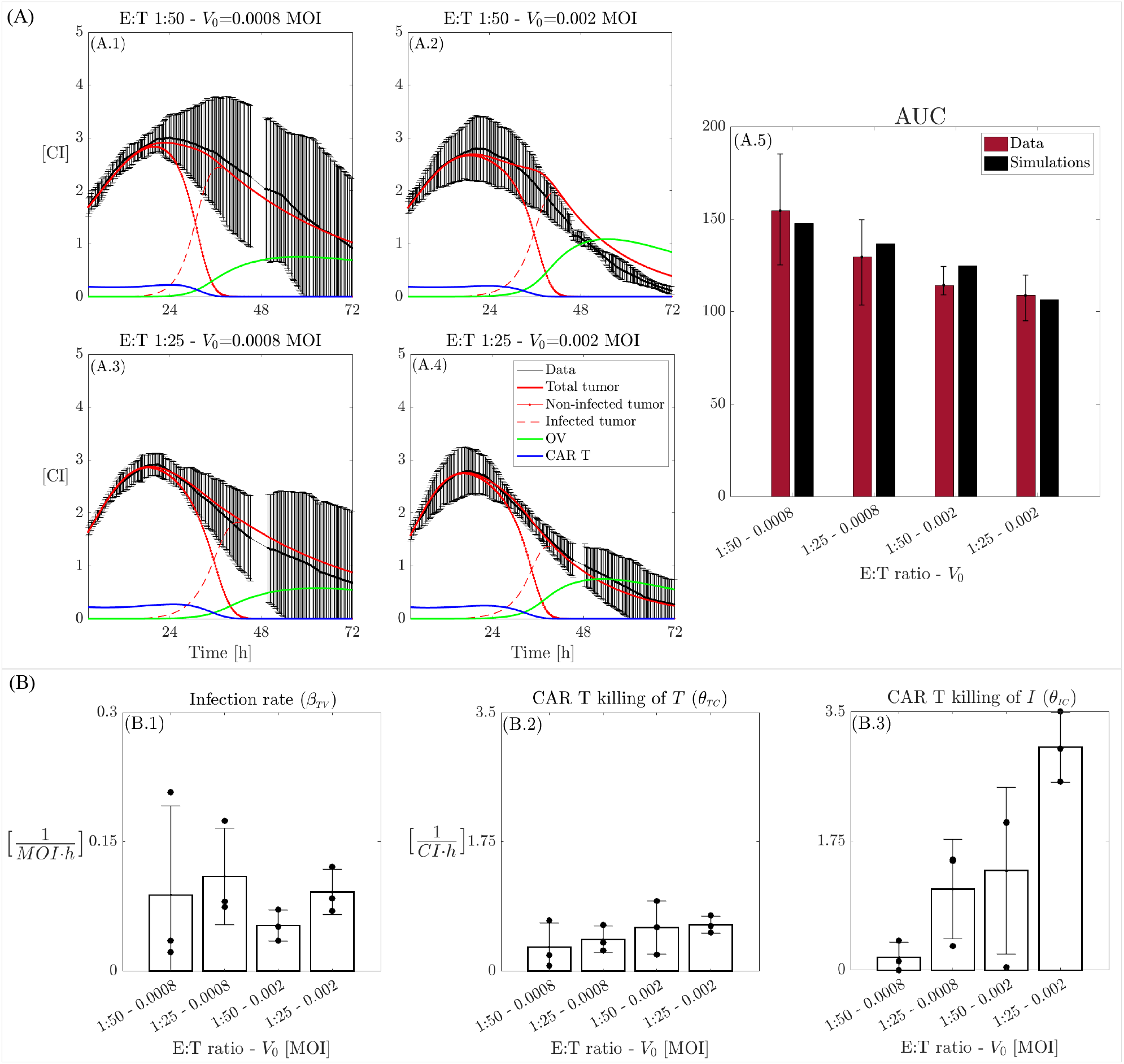
Dynamics of model (1) and *in vitro* CAR T-cell, OV, and glioma cell data. A: Dynamics of cancer cells along with the model fits (black: data, red continuous line: total tumor cells, red dashed–dot line: non-infected tumor cells, red dashed line: infected tumor cells, green line: OV, blue line: CAR T-cells) with initial E:T ratio and virus dose of (A.1) 1:50–0.0008 MOI, (A.2) 1:50–0.002 MOI, (A.3) 1:25–0.0008 MOI, and (A.4) 1:25–0.002 MOI. (A.5) Comparison of AUC obtained from the *in vitro* data (red) and *in silico* (black) simulations. (B) Tumor infection rate (*β*_*TV*_), CAR T killing rate of tumor cells (*θ*_*TC*_), and CAR T killing rate of infected tumor cells (*θ*_*IC*_). Each parameter is plotted for the four different combinations of effector to target ratios and initial virus concentrations (1:50–0.0008, 1:25–0.0008, 1:50–0.002, and 1:25–0.002). Bars represent the mean values of the parameters, while error bars indicate the standard deviation. Black bullets denote the individual values for each of the three replicates from the experiments.

To analyze the effects of the therapy combination compared to the two monotherapy cases, we directly compare the characteristic parameters of each monotherapy: the infection rate *β*_*TV*_ and the CAR T killing rate of non-infected tumor cells *θ*_*TC*_. This comparison is made for two initial virus concentrations, *V*_0_ = 0.0008 and *V*_0_ = 0.002, as well as for two E:T ratios, 1:50 and 1:25 –the doses tested in both the monotherapy and combination scenarios. Additionally, we assess the value of AUC for the corresponding monotherapy and combined therapy cases. The results of this analysis are presented in Figure 9. Comparing the infection rate *β*_*TV*_ estimated for the OV monotherapy and CAR T–OV therapy, there is a clear increase in the effectiveness of OV infection in the combined-therapy case with respect to the monotherapy scenario. Although the differences are less pronounced, there is also a noticeable trend indicating that CAR T–cell efficacy in killing tumor cells increases in the combined-therapy scenario compared to monotherapy [36, 50], especially for the initial E:T ratio 1:25. Figure 9–E clearly shows that the combined therapy is more effective in reducing the AUC compared to the individual treatments. In the monotherapy setting, increasing the E:T ratio from 1:50 to 1:25 does not significantly affect the AUC, however, raising *V*_0_ from 0.0008 to 0.002 substantially reduces AUC. Overall, the combination therapy consistently demonstrates better outcomes across all scenarios, with improvements observed as either *V*_0_ or the initial E:T ratio increases.

**Figure 9:**
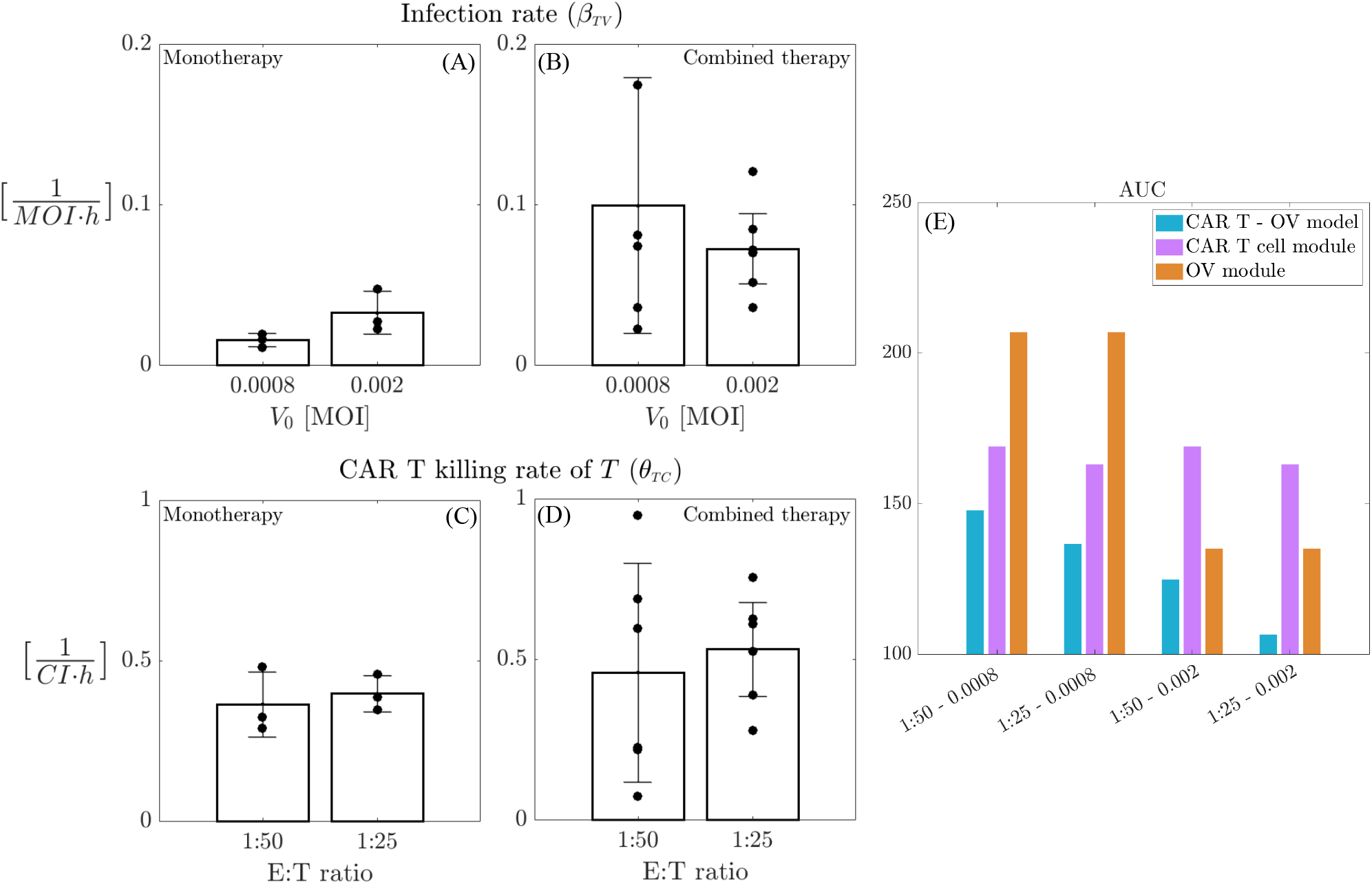
Benefits of combined therapies. Comparison of parameter estimations for monotherapy and combined therapy scenarios. A-B: comparison of *β*_*TV*_ values for initial OV concentration of *V*_0_ = 0.0008 and *V*_0_ = 0.002 in the monotherapy (A) and combined therapy (B) cases. C-D: comparison of *θ*_*TC*_ values for initial E:T ratios 1:50 and 1:25 in the monotherapy (C) and combined therapy (D) cases. Panel E compare the *in silico* AUC values for the monotherapy (purple bars for E:T of 1:50 and 1:25, orange bars for *V*_0_ = 0.0008 and *V*_0_ = 0.002) and combined therapy (light–blue bars) cases. Corresponding therapy doses for each case (mono– or combined therapy) are indicated below each group.

#### 3.3.2 CAR T–OV model predicts tumor outcomes for different therapeutic combinations

We evaluated our model’s ability to predict tumor outcomes from combinations of therapy doses that differ from those in the analyzed dataset. Experimentally, several challenges may arise when managing and utilizing very low concentrations of therapeutic agents. Specifically, experiments involving low doses of viruses often exhibit significant stochastic variability, leading to less reliable results. Nevertheless, lower therapy doses are advantageous for patient treatment, as they can minimize potential side effects. In this context, *in silico* experiments serve as powerful tools for testing hypotheses about therapy combinations and offer valuable insights into potential therapeutic outcomes. To this end, we analyze how our model responds to four additional combinations of CAR T-cells and oncolytic virus. We maintain the same E:T ratios as in Section 3.3.1, specifically 1:50 and 1:25, but assume lower initial virus concentrations, with *V*_0_ = 6 ×10^−4^, 1.2 ×10^−5^ MOI. Assuming that we do not have access to the data about the temporal evolution of the tumor population in these scenarios, we use AUC as a measure of treatment outcome.

We apply model (1) to estimate AUC and compare the *in silico* results with the *in vitro* data. Instead of fitting the model to the data, we leverage the parameter space conclusions from previous sections to estimate parameters for these scenarios. Specifically, we assume that parameters *µ* and *δ* remain unaffected by oncolytic virus particles, using the corresponding estimates obtained in Section 3.1. Parameters *b, ω*, and *K* are fixed as shown in Figure 7. For the remaining parameters, we utilize an interpolation procedure. For each initial E:T ratio, we estimate *α, β*_*TV*_, *β*_*CV*_, *θ*_*TC*_, and *θ*_*IC*_ using the results from model (1) for the two different initial virus concentrations. We apply polynomial interpolation to derive the corresponding parameter values for the new *V*_0_ concentrations of 6 ×10^−4^ and 1.2 ×10^−5^ MOI. A similar procedure is employed to determine the new value of the parameter *γ*, based on the estimates from model (3). The results of this analysis are presented in Figure 10, showcasing the comparison between AUC obtained from *in vitro* experiments (red) and the numerical simulations (black). From Figure 10, we observe that our model demonstrates strong predictive capability, providing accurate estimates for AUC, particularly at *V*_0_ = 6 ×10^−4^. Additionally, a comparison with Figure 8-A.5 reveals an interesting trend: for lower E:T ratios, similar quantitative values for AUC can be achieved by decreasing *V*_0_ from 8 ×10^−4^ to 6 ×10^−4^ and also from 6 ×10^−4^ to 1.2 ×10^−5^. Our results suggest that the synergy between the two treatments supports the use of virus particles at lower initial concentrations, without compromising tumor cell kill. We provide the qualitative evolution of the model in the four different scenarios in Supplementary Figure S.5 the Supplementary Material S.4.

**Figure 10:**
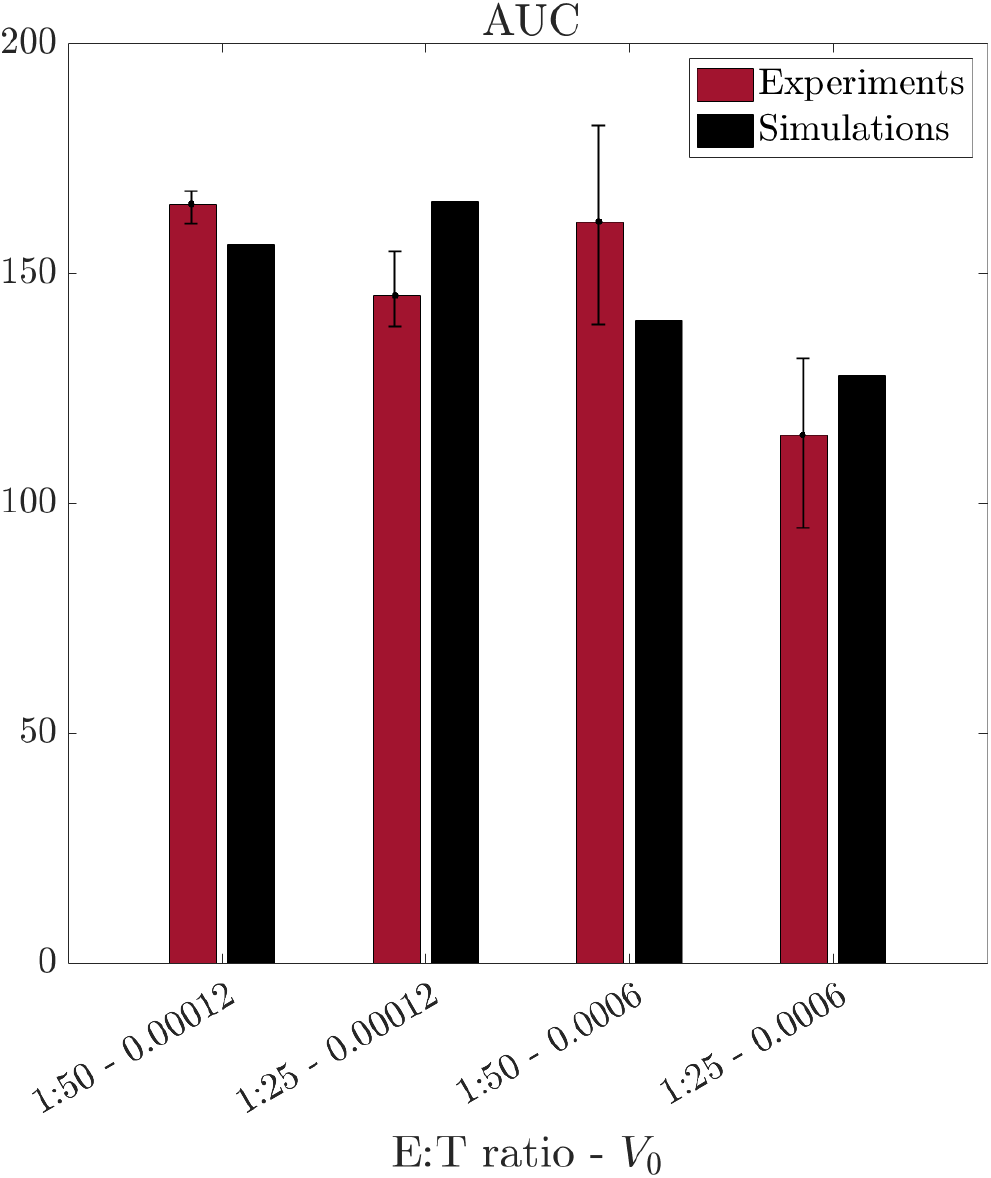
Prediction of tumor response to different therapy combinations. Comparison between the values of AUC obtained from the *in vitro* (red) and *in silico* (black) data. The numerical simulations are conducted without data fitting, instead predicting tumor behavior for therapy combinations that differ from those in the described datasets.

#### 3.3.3 Opportunity window for therapy administration

Finally, we utilize our developed framework to predict potential tumor outcomes based on different therapy administration schedules. Specifically, we analyze the effects of delaying the administration of one or both therapies within a time window of [0-40] hours. We measure the impact of these varying schedules on AUC as well as on the tumor population 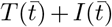 at the final time point 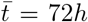. The findings of this analysis are presented in Figure 11.

**Figure 11:**
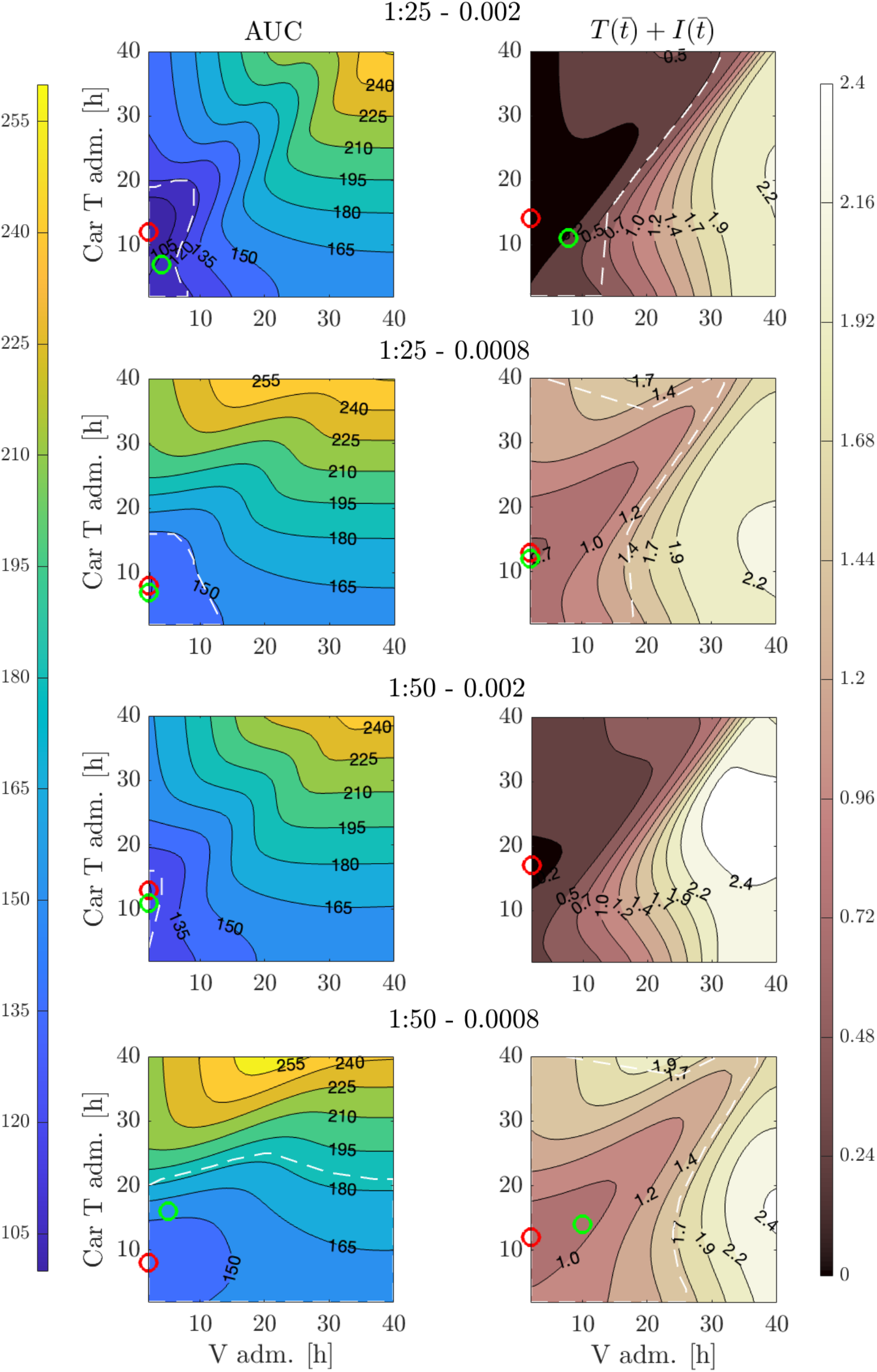
Effects of timing in combined therapy administration. Contour plots illustrate AUC (left column) and total tumor population 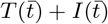 at the final time point 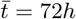 (right column) calculated from model (1) for various therapy administration schedules. The *x*–axis represents the time of oncolytic virus administration, while the *y*–axis represents the timing of CAR T-cell administration. Each row corresponds to a different combination of therapy doses: E:T=1:25 and *V*_0_ = 0.002 (first row); E:T=1:25 and *V*_0_ = 0.0008 (second row); E:T=1:50 and *V*_0_ = 0.002 (third row); E:T=1:50 and *V*_0_ = 0.0008 (fourth row). In each subplot, the red marker indicates the combination of therapy delays that minimizes the illustrated quantity,while the green marker indicates the combination that yields a value equal to the experimental mean for the same quantity. White dashed lines outline the region defined by the experimental standard deviation of the corresponding quantity.

For each therapy combination, we present the variety of possible tumor outcomes that can arise from delaying the two therapeutic treatments. Specifically, we use a red marker to indicate the combination of therapy delays that yields the best tumor outcome, characterized by the lowest AUC (left column) or the lowest final tumor population (right column). Additionally, based on the mean and standard deviation values obtained from the experiments, we employ a green marker to denote the combination of therapy delays that leads to a AUC value (left column) or final tumor population value (right column) corresponding to the experimental mean. The white dashed lines outline the confidence region in the administration schedule space defined by the standard deviation of the experimentally measured outcomes.

Our analysis reveals an interesting nonlinearity in the effects of therapy delays concerning the administration timing of both OV and CAR T-cells across all cases. Notably, greater changes in tumor outcomes, both in terms of AUC and 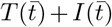, result from delays in CAR T-cell administration, while variations in OV administration yield smaller effects. This observation may have a biological explanation: delaying CAR T cell administration allows more time for the virus to infect tumor cells, potentially weakening them and making them easier for CAR T-cells to eliminate [50]. Additionally, this delay may help reduce CAR T cell exhaustion, thereby enhancing the overall efficiency of the immune response against the tumor. Furthermore, delays in CAR T-cell administration are crucial for achieving the best improvements in tumor outcomes, as evidenced by the minimum values for both measurements (red markers). With the exception of 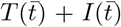 for the combination 1:50 – 0.002, all other cases demonstrate that delays in therapy lead to improved outcomes, indicating that the achieved minima are lower than the experimental values obtained when therapies are administered together at the initial time. For the combination of 1:50 – 0.002, no improvements were observed with delays in either treatment. This may indicate that the OV concentration is sufficiently high relative to the E:T ratio, suggesting that delaying administration may not enhance treatment outcomes. However, it also implies that further data from the experiment could provide additional insights to better understand this prediction. The regions outlined by the white dashed lines indicate where the treatment results fall within the experimental error bars, suggesting some flexibility in the therapy schedule without compromising final tumor outcomes. Finally, we compare the combined therapy results with those from monotherapy, which are summarized in the Supplementary Figure S.6 the Supplementary Material S.4. In all scenarios, the impact of therapy delays in the monotherapy cases is significantly less than that observed in the combined treatment, and no monotherapy scenario leads to an improvement in overall tumor outcomes when delay in the therapy administration time is considered.

## 4 Discussion

We have developed a mathematical model to describe the complex dynamics involved in glioblastoma cancer cell growth, CAR T-cell killing and proliferation, and oncolytic virus infection. The mathematical model was constructed with the principle of parsimony in mind, incorporating only the most essential variables and the simplest assumptions necessary to capture the key interactions of the system. Leveraging knowledge from the fields of ecology and epidemiology, we integrated a predator-prey model to represent the dynamics between cancer cells and CAR T-cells, and a SIR-type model to describe the viral infection process and resulting CAR T-cell killing. This combination enabled us to investigate the synergistic effects of these therapies when applied together. The model was fit to experimental data, leading to several insights and observations.

A significant finding of our modeling and analysis was the impact of virus-specific parameters, particularly the bursting size *b*, on the dynamics of the system. This is because the burst size is logarithmically related to the infection rate, meaning that it can significantly influence viral load and, in turn, the therapeutic efficacy against tumor cells. This observation is important because values of virus burst size may vary by orders of magnitude in the literature [19], and—more importantly—it is effectively *unmeasurable* in *in vivo* settings. The critical role of the burst size is evident through the model’s predictions: if the burst size is too high, it may lead to viral saturation, reducing viral efficacy and impairing CAR T-cell function. Conversely, if the burst size is very low, the infection dynamics become more stochastic than deterministic and make prediction of therapeutic efficacy difficult [43]. We therefore conducted an analysis of how the bursting size affects various therapeutic scenarios and found that identifying an optimal burst size plays a pivotal role in balancing the therapeutic effects of the virus and CAR T-cells, paving the way for improved therapeutic outcomes.

Another insight from our modeling was the identification of optimal treatment timing for the combined therapies. Through simulations, we explored the impact of therapy scheduling on two key metrics of therapeutic efficacy: the AUC and the final number of tumor cells. We discovered non-linear relationships between the timing of oncolytic virus administration and CAR T-cell therapy. Specifically, the most effective outcomes consistently occurred when the OV was administered simultaneously with, or prior to, the CAR T-cells. Delaying OV administration by more than 24 hours after CAR T-cell infusion significantly diminished its therapeutic benefit. This finding highlights the importance of timing in combination therapies and suggests the most effective treatment regimens. These insights could lead to more optimized treatment strategies, particularly in the context of multi-dose regimens involving either OV or CAR T-cells [50].

It is important to note that our experiments, modeling, and analysis are based on a simplified *in vitro* system, which does not account for important *in vivo* factors such as host immunity, the tumor microenvironment, or other potential modulators of OV and CAR T-cell activity. While these factors are undoubtedly critical in realworld therapeutic settings, the simplified experimental setup and mathematical modeling approach we have employed represent a crucial first step in understanding the complex, nonlinear dynamics underlying these two therapies. This work lays the foundation for future research that will incorporate more complex *in vivo* factors and refine therapeutic strategies in the pursuit of more effective cancer treatments.

## Acknowledgments

Research reported in this work was supported by the National Cancer Institute of the National Institutes of Health under grant numbers R01CA254271 (C.E.B.), R01NS115971 and P30CA033572 (C.E.B., R.C.R.), and the Marcus Foundation, the California Institute of Regenerative Medicine (CIRM) under CLIN2-10248, and a sponsored research agreement from Mustang Bio (C.E.B.). M.C. acknowledges support from City of Hope’s Global Scholar Program. M.C. was supported by the National Group of Mathematical Physics (GNFM-INdAM) through the INdAM–GNFM Project (CUP E53C22001930001) *From kinetic to macroscopic models for tumor-immune system competition*, and by the European Union - NextGenerationEU and the MUR-Italian Ministry of Universities and Research through the project PRIN 2022 PNRR *Mathematical modeling for a Sustainable Circular Economy in Ecosystems* (project code P2022PSMT7, CUP D53D23018960001).

## Data availability statement

All relevant data are within the manuscript and its Supporting Information files. The raw data and the Matlab codes are available in the GitHub repository https://github.com/mconte93/CAR-OV-study.git.

## Author contribution statement

Conceptualization: A.X., M.C., C.E.B., R.C.R., Methodology: M.C., A.X., R.T.W., S.B., R.C.R., Formal analysis: M.C., A.X., R.T.W., R.C.R., Resources: C.E.B., K.A.C., R.C.R., Writing—original draft preparation: M.C., A.X., R.T.W., R.C.R., Writing—review and editing: M.C., A.X., R.T.W., S.B., K.A.C., C.E.B., R.C.R., Supervision: R.C.R., Project administration: C.E.B., R.C.R., Funding acquisition: C.E.B., K.A.C., R.C.R., All authors contributed to the article and approved the submitted version.

## List of Supplementary Information

**S.1 Qualitative analysis of the dynamical systems**

**S.2 Parameter value estimations**

**S.3 Extended analysis of burst size**

**S.4 Extended predictions of CAR T–OV model**

**Supplementary Figure S.1**: **Analysis of the dynamical system** (S.2). Phase space diagrams of the non-dimensionalized **CAR T module** given by equations (S.2) in three different scenarios: (A) *A > B*, with two equilibrium points; (B) *A < B*, with three equilibrium points; (C) *A* ≈ 0 and *B >* 0, with equilibrium *E*_2_ and an infinite set of equilibria *E*_*N*_ with *Y*_1_ = 0. In (D) we show the bifurcation diagram of *Y*_1_ with respect to the parameter *B*. Parameter values are set to *A* = 0.58 and *B* = 0.026 (A), *A* = 0.58 and *B* = 1.31 (B), and *A* = 0 and *B* = 0.026 (C). The figure is embedded in the Section S.1.1.

**Supplementary Figure S.2**: **Analysis of the dynamical system** (S.5). Phase space diagrams of the non-dimensionalized **OV module** given by equations (S.5) in three different scenarios: (A) *E < Ē*, with two equilibrium points; (B) 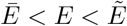, with three equilibrium points; (C) 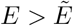, where the limit cycle emerges. In (D) we show the bifurcation diagram for *Y*_1_ with respect to the parameter *E*. Parameter values are set to *D* = 0.58, *F* = 0.48, and *E* = 0.15 (A), *E* = 1 (B), or *E* = 3.5 (C). The figure is embedded in the Section S.1.2.

**Supplementary Figure S.3**: **Burst size effects on the tumor growth and apoptosis rates**. Estimation of the parameters *α* (first row) and *γ* (second row) from the **OV module** given by system (S.4) for various burst size values *b* ∈ [0.025, 2500]. Columns correspond to the three different initial virus doses: *V*_0_ = 0.03, *V*_0_ = 0.002, and *V*_0_ = 0.0008. The figure is embedded in the Section 3.2.1.

**Supplementary Figure S.4**: **Infected cell fraction analysis**. (A): Black circles represent experimental data for the fraction of infected tumor cells (*I*(*t*)) relative to the total tumor population (*I*(*t*) + *T* (*t*)) at 24 hours (left) and 48 hours (right) for virus doses *V*_0_ ∈ {0.0005, 0.001, 0.01, 0.1, 1}. Red stars indicate the fractions of infected tumor cells at 24 hours (left) and 48 hours (right) for virus doses *V*_0_ ∈ {0.0008, 0.002, 0.03}, derived through interpolation. Initial virus concentrations are presented on a log scale. Data are shown with means (markers) and standard deviations (error bars). (B): Contour plots depicting the fraction of infected tumor cells (*I*(*t*)) relative to the total tumor population (*I*(*t*) + *T* (*t*)) at 24 hours (top row) and 48 hours (bottom row). The three columns correspond to initial virus concentrations *V*_0_ = 0.0008, 0.002, 0.03, respectively. The fraction of infected cells is illustrated as a function of the virus clearance rate *ω* (x–axis) and the burst size *b* (y–axis, in log scale). In each subplot, the mean value of the interpolated experimental data (red markers in panel A) is represented by a red solid line, while the standard deviation (red error bars in panel A) is shown with a red dashed line. The figure is embedded in the Section 3.2.1.

**Supplementary Figure S.5**: **Dynamics of model** (1) **and** *in vitro* **CAR T-cell, OV, and glioma cell data**. Dynamics of cancer cells, expressed in cell index from xCELLigence, along with the model trajectories obtained from system (1) (black: data; red continuous line: total tumor cells; red dashed–dot line: non-infected tumor cells; red dashed line: infected tumor cells; green line: OV; blue line: CAR T-cells) with initial E:T ratio and virus dose of 1:25–0.0006 MOI (A), 1:50–0.0006 MOI (B), 1:25–0.00012 MOI (C), and 1:50–0.00012 MOI (D). The parameters used in each of the four combinations of therapy are described in Section 3.3.2 in the main text. The figure is embedded in the Section S.4.

**Supplementary Figure S.6**: **Effects of delay in monotherapy administration**. (A): CAR T-cell monotherapy case. AUC (left column) and tumor population at the final time point 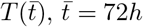, (right column) calculated from model (S.1) for different therapy administration schedule. The *x*–axis represents the time of CAR T cell administration. In each plot the effect of therapy delay is shown for E:T=1:50 (continuous line), E:T=1:25 (dashed line), and E:T=1:10 (dot-dashed line). (B): OV monotherapy case. AUC (left column) and tumor population at the final time point 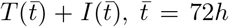, (right column) calculated from model (S.4) for different therapy administration schedule. The *x*–axis represents the time of OV administration. In each plot the effect of therapy delay is shown for *V*_0_ = 0.0008 MOI (continuous line),*V*_0_ = 0.002 MOI (dashed line), and *V*_0_ = 0.03 MOI (dot-dashed line). The figure is embedded in the Section S.4.

**Supplementary Table S.1**: **Parameter fitting for model** (S.1). Average values of the parameters *α, θ*_*TC*_, *µ*, and *δ* from model (S.1), obtained from the fitting to the CAR T-cell and glioma cell data. The table is embedded in the Section S.2.

**Supplementary Table S.2**: **Parameter fitting for model** (S.4). Average values of the parameter *α* and *γ* from model (S.4), obtained from the fitting to the OV and glioma cell data for burst size values *b* ∈ [0.025, 2500] MOI/CI. The remaining parameter are fixed to *K* = 5.2246 CI and *ω* = 0.05 *h*^−1^. Estimations of *β*_*TV*_ are reported in Table S.4 and in Figure 5 in the main text. The table is embedded in the Section S.2.

**Supplementary Table S.3**: **Parameter fitting for model** (1). Average values of the parameter *α, β*_*TV*_, *θ*_*TC*_, *θ*_*IC*_, and *β*_*CV*_ from model (1), obtained by fitting to the CAR T-cells, OV, and glioma cell data. The remaining parameter are fixed to *K* = 5.2246 CI, *b* = 25 MOI/CI, *ω* = 0.05 *h*^−1^. *γ* is taken from the corresponding estimation in Table S.2, while *δ* and *µ* are taken from the corresponding estimations in Table S.1. The table is embedded in the Section S.2.

**Supplementary Table S.4**: **Effects of burst size variation on model** (S.4) **parameters**. Values of the parameters *α, β*, and *γ* of model (S.4) for different values of the burst size *b* ∈ [0.025, 2500] MOI/CI. The table is embedded in the Section 3.2.1.

