## Supplementary Information S1 for "CAR T-cell and oncolytic virus dynamics and determinants of combination therapy success for glioblastoma"

### S.1 Qualitative analysis of the dynamical systems

Here we present some results that complement and complete the analysis presented in the main text. Precisely, we consider the two dynamical systems given by equations (S.1) and (S.4) and we summarize their characteristics in terms of equilibria, stability, and bifurcations.

#### S.1.1 CAR-T cell module

For describing the interactions between tumor cells ($T$) and CAR T cells ($C$), we consider the following system of ODEs

$$\begin{cases} \dot{T} = \alpha T\left(1 - \dfrac{T}{K}\right) - \theta_{TC} TC \\ \dot{C} = -\delta C + \mu CT\,. \end{cases} \tag{S.1}$$

This model employs classical predator–prey dynamics to investigate the mechanisms of CAR T-cell killing in GBMs. Here, tumor cells are proliferating at rate $\alpha$ up to a carrying capacity $K$. CAR T cells kill tumor cells at rate $\theta_{TC}$, while they naturally die at rate $\delta$. All these parameter are assumed to be positive. Moreover, we include a mechanism of CAR T cells stimulation/exhaustion by tumor cells at rate $\mu$, which can take either positive or negative values. Rescaling the temporal variable with respect to the characteristic time of tumor cell proliferation, i.e., setting $\tau = \alpha t$, we can non-dimensionalized the system by setting $Y_1 = T/K$, and $Y_4 = (\theta_{TC}/\alpha)C$. System (S.1) now reads

$$\begin{cases} \dfrac{dY_1}{d\tau} = Y_1\,(1 - Y_1) - Y_1 Y_4 \\ \dfrac{dY_4}{d\tau} = -AY_4 + B\,Y_1 Y_4\,. \end{cases} \tag{S.2}$$

where $A = \delta/\alpha$ is the normalized death rate of CAR T cells, while $B = (K/\alpha)\mu$ is the normalized killing rate of tumor cells. For this system, it is easy to prove that the first quadrant is a positively invariant set, ensuring the positivity of solutions when starting from positive initial conditions. Additionally, the trajectories $Y_1(t) = 0$ and $Y_4(t) = 0$, $\forall t > 0$, cannot be crossed.

The system has three possible equilibrium configurations given by

$$E_1 = (0, 0)\ , \qquad E_2 = (1, 0)\ , \qquad E_3 = \left(\frac{A}{B}, \frac{B - A}{B}\right)$$

where $E_3$ belong to the phase plane only for $B \geq A$. In particular, $E_3$ reflects a coexistence scenario. Biologically this condition of the existence of $E_3$ means that a coexistence equilibrium between tumor cells and CAR T cells is possible only when CAR T cells are enhanced by tumor cell (i.e., $\mu > 0$) and they are strong enough to fight the tumor cells. Local stability properties of the equilibrium states can be determined by analyzing the eigenvalues of the Jacobian matrix associated with the system (S.2). The Jacobian at $E_1$, denoted as $\mathcal{J}(E_1)$, is a diagonal matrix with diagonal elements 1 and $-A$, i.e., $E_1$ is always unstable. Instead $\mathcal{J}(E_2)$ is an upper triangular matrix with diagonal elements $\lambda_1 = -1$ and $\lambda_2 = B - A$. Therefore, $E_2$ is locally asymptotically stable if and only if $A > B$. Thus, when $E_3$ exists, $E_2$ is always unstable. Finally, for the stability of $E_3$, the Jacobian matrix $\mathcal{J}(E_3)$ is given by

$$\mathcal{J}(E_2) = \begin{pmatrix} -\dfrac{A}{B} & -\dfrac{A}{B} \\ B - A & 0 \end{pmatrix}\,. \tag{S.3}$$

In the region where $E_3$ is an admissible equilibrium, distinct from $E_2$ (i.e., for $B > A$), the determinant and trace of the Jacobian are

$$det(\mathcal{J}(E_3)) = \frac{A}{B}(B - A) > 0 \qquad \text{and} \qquad Tr(\mathcal{J}(E_3)) = -\frac{A}{B} < 0\,.$$

This means that $E_3$ is always stable. At $A = B$, there is a transcritical bifurcation of the forward type. In particular, the condition $A = B$, which enables CAR T cells to coexists with tumor cells, relates to the effectiveness of immune cells, which is enhanced by the tumor cells, compensating CAR T natural death. In any case, the treatment is only able to control tumor grow, with tumor cell density as small as the fraction A/B, but it cannot eradicate the cancer cells. As observed in [1], a successful CAR T-cell treatment can be obtained only by assuming that CAR T cells death is negligible, i.e., $A \approx 0$. In this case, besides $E_2$, we obtain an infinite set of equilibrium points characterized by $Y_1 = 0$. Here, $E_2$ is locally asymptotically stable if and only if $B < 0$, namely for CAR T cell exhaustion, while the equilibrium points $E_N = (0, Y_4)$ can be locally stable when $Y_4 \geq 1$, i.e., if the net rate of CAR T cell killing ($C\theta_{TC}$) is greater than the proliferation rate of the cancer cells. For completeness, in Figure S.1 we show the phase space diagrams related to system (S.2) for $A > B$ (panel A) and $A < B$ (panel B), as well as, when $A \approx 0$ and $B > 0$ (panel C). Moreover, we include the qualitative bifurcation diagram related $Y_1$ and the parameter $B$ (panel D). By examining the values of $A$ and

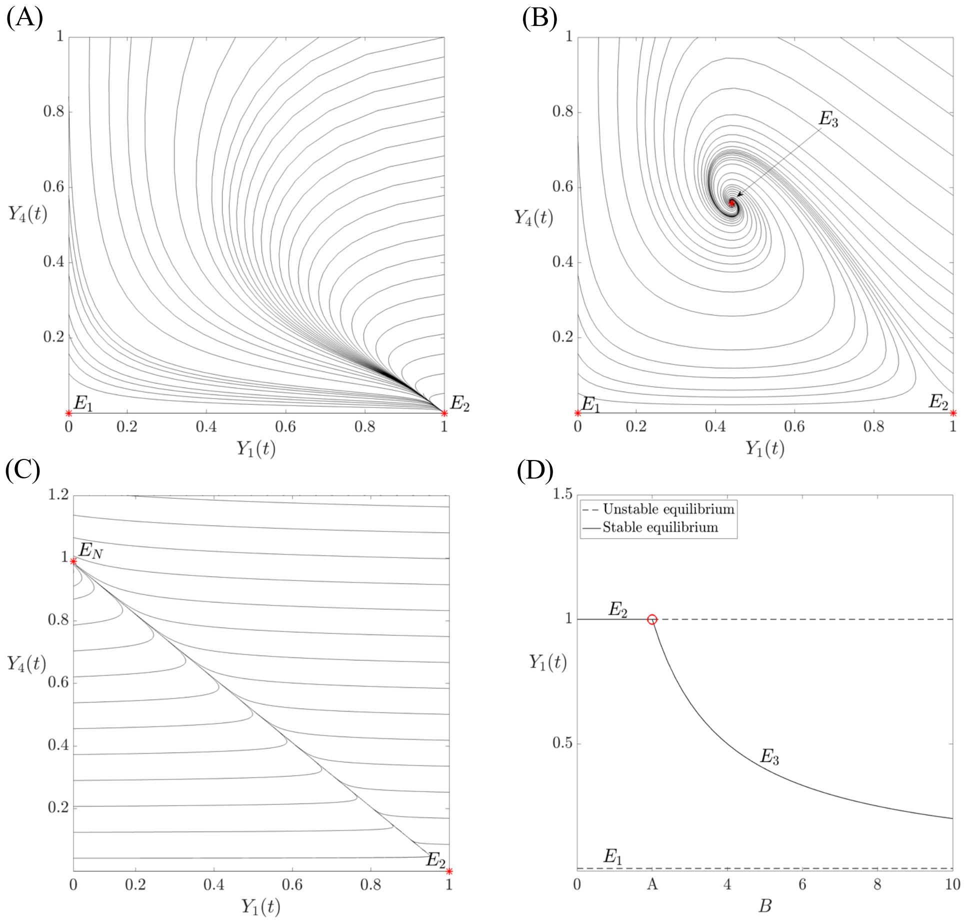


Supplementary Figure S.1: **Analysis of the dynamical system** (S.2). Phase space diagrams of the non-dimensionalized **CAR T module** given by equations (S.2) in three different scenarios: (A) $A > B$, with two equilibrium points; (B) $A < B$, with three equilibrium points; (C) $A \approx 0$ and $B > 0$, with equilibrium $E_2$ and an infinite set of equilibria $E_N$ with $Y_1 = 0$. In (D) we show the bifurcation diagram of $Y_1$ with respect to the parameter $B$. Parameter values are set to $A = 0.58$ and $B = 0.026$ (A), $A = 0.58$ and $B = 1.31$ (B), and $A = 0$ and $B = 0.026$ (C).

$B$, which can be derived from the parameter values obtained through data fitting in Section 3.1, we observe that for all three E:T ratios, the parameters satisfy $A < B$. This implies that, given enough time, the model will asymptotically approach the equilibrium state $E_3$.

##### S.1.2 OV module

For describing the interactions between tumor cells ($T$), infected tumor cells ($I$), and oncolytic virus ($V$) we consider the following system of ODEs

$$\begin{cases} \dot{T} = \alpha T \left(1 - \dfrac{T+I}{K}\right) - \beta_{TV} TV \\ \dot{I} = \beta_{TV} TV - \gamma I \\ \dot{V} = \gamma b I - \omega V\,. \end{cases} \tag{S.4}$$

This model employs classical SIR dynamics to investigate the mechanisms of tumor infection due to the OV particles and consequent cell apoptosis. Here, non-infected tumor cells are proliferating at rate $\alpha$ up to a carrying capacity $K$. They are infected due to interactions with the virus particles at rate $\beta_{TV}$ and the infected tumor cells undergo apoptosis at rate $\gamma$. This process determine the release of new virus particles in the environment at a rate proportional to $\gamma$, with proportionality constant $b$, which represent the virus burst size. Virus particles are also cleared from the system at rate $\omega$. Rescaling the temporal variable with respect to the characteristic time of tumor cell proliferation, i.e., setting $\tau = \alpha t$, we can non-dimensionalized the system by setting $Y_1 = T/K$, $Y_2 = I/K$, and $Y_3 = (\beta/\alpha)V$. System (S.4) now reads

$$\begin{cases} \dfrac{dY_1}{d\tau} = Y_1\left(1 - Y_1 - Y_2\right) - Y_1 Y_3 \\ \dfrac{dY_2}{d\tau} = Y_1 Y_3 - D Y_2 \\ \dfrac{dY_3}{d\tau} = E Y_2 - F Y_3 \end{cases} \tag{S.5}$$

where $D = \gamma/\alpha$ is the rescaled apoptosis rate of infected tumor cells, while $E = (\beta\gamma\, b\, K)/\alpha^2$ and $F = \omega/\alpha$ are the rescaled production and cleareance rate of the virus, respectively. For this system, it is easy to prove that the first octant is a positively invariant set, ensuring the positivity of solutions when starting from positive initial conditions. Additionally, the plane $Y_1(t) = 0$, $\forall t > 0$, cannot be crossed.

The system has three possible equilibrium configurations given by

$$E_1 = (0,0,0)\ , \qquad E_2 = (1,0,0)\ , \qquad E_3 = \left(\frac{DF}{E}, \frac{F}{E}\frac{E-DF}{F+E}, \frac{E-DF}{F+E}\right)$$

where $E_3$ belong to the phase plane only for $E \geq DF$. In particular, $E_3$ reflects a coexistence scenario whose conditions for existence biologically can be translating by requiring the virus production by bursting to happen faster than the virus clearance. Local stability properties of the equilibrium states can be determined by analyzing the eigenvalues of the Jacobian matrix associated with the system (S.5). The Jacobian at $E_1$ is a diagonal matrix with diagonal elements 1, $-D$, and $-F$ i.e., $E_1$ is always unstable. Instead, $\mathcal{J}(E_2)$ is given by

$$\mathcal{J}(E_2) = \begin{pmatrix} -1 & 0 & 0 \\ 0 & -D & 1 \\ 0 & E & -F \end{pmatrix}, \tag{S.6}$$

where the eigenvalue $\lambda_1 = -1$ is immediately apparent. The signs of the remaining two eigenvalues can be determined from the determinant and trace of the submatrix obtained by deleting the first row and first column. Using this, we can infer that $E_2$ is locally asymptotically stable if and only if $DF > E$. Thus, when $E_3$ exists, $E_2$ is always unstable. At $E = \bar{E} = DF$ there is a transcritical bifurcation of the forward type. Finally, for the stability of $E_3$, the Jacobian matrix $\mathcal{J}(E_3)$ is given by

$$\mathcal{J}(E_3) = \begin{pmatrix} -Y_1^* & -Y_1^* & -Y_1^* \\ \dfrac{E(1-Y_1^*)}{E+F} & -D & Y_1^* \\ 0 & E & -F \end{pmatrix}, \tag{S.7}$$

where we indicate with $Y_1^* := (DF)/E$. The characteristic equation of $\mathcal{J}(E_3)$ is given by $\lambda^3 + P_2\lambda^2 + P_1\lambda + P_0 = 0$, where the coefficients $P_i$, for $i = 0, 1, 2$, are given by

$$P_2 = Y_1^* + F + D$$

$$P_1 = Y_1^*\left(F + D + \frac{E(1-Y_1^*)}{E+F}\right)$$

$$P_0 = E Y_1^*(1 - Y_1^*)$$

Using the Routh-Hurwitz criterion, we obtain that the equilibrium point $E_3$ is locally asymptotically stable if $E \geq DF$ (condition of existence) and it holds $P_2 > 0$, $P_1 > 0$, and $P_1 P_2 - P_0 > 0$. While the first two conditions are always true, we need to study in which regions of the parameter space $P_1 P_2 - P_0 > 0$. We get

$$\begin{aligned} P_1 P_2 - P_0 &= \frac{DF}{E^2(E+F)}\left(-E^3 + E^2((F+D)^2 + D(F+1)) + FE((F+D)^2 + D(F+1)) + F^3 D\right) \\ &= \frac{DF}{E^2(E+F)}\mathcal{H}(E) := \mathbf{H}(E)\,. \end{aligned}$$

The sign depends on $\mathcal{H}(E)$, which has 2 negative roots and 1 positive roots. As we are interested in the positive values for $E$, there exists a unique $\tilde{E} > 0$ such that $\mathcal{H}(\tilde{E}) = 0$. Moreover, for $E \to \infty$ we observe that $\mathcal{H} \to -\infty$, i.e., $\mathcal{H} > 0$ for $E < \tilde{E}$ and $\mathcal{H} < 0$ for $E > \tilde{E}$. Calculating $\mathcal{H}(\bar{E})$, we notice that $\mathcal{H}(\bar{E}) > 0$, thus we can conclude that $\bar{E} < \tilde{E}$ and that for $\bar{E} < E < \tilde{E}$ the equilibrium point $E_3$ is locally asymptotically stable. Using the Liu's criterion [2], we can show that at $E = \tilde{E}$ an Hopf bifurcation occurs. In fact, $\mathbf{H}(\tilde{E}) = 0$ and

$$\frac{d\mathbf{H}}{dE}|_{E=\tilde{E}} = -\frac{DF}{E^3(E+F)}\left((m+F)E^3 + 2mFE^2 + (F^2m + 3F^3D)E + 2F^4D\right)|_{E=\tilde{E}} \neq 0 \qquad \forall E > 0$$

since all the coefficients (with respect to $E$) are positive, with $m := (F+D)^2 + D(F+1)$. These two conditions satisfy the Hopf bifurcation requirements. These results are perfectly in line with the one obtained in [3]. Overall, the results suggest that for obtaining a stable scenario of tumor growth under control, the virus production through cell bursting must balance virus clearance, (i.e., $E \geq \bar{E}$). However, if virus production becomes too pronounced relative to other processes, such that $E \geq \tilde{E}$, the system may exhibit oscillatory behavior, potentially leading to the emergence of a limit cycle. These findings also highlight the crucial role of virus-specific parameters, particularly $b$ and $\omega$, in determining the outcome of therapeutic interventions. For completeness, in Figure S.2 we show the phase space diagrams related to system (S.5) for $E < \bar{E}$ (panel A), $\bar{E} < E < \tilde{E}$ (panel B), and $E > \tilde{E}$ (panel C). Moreover, we include the qualitative bifurcation diagram related $Y_1$ and the parameter $E$ (panel D). By examining the values of $D$, $E$, and $F$, which can be derived

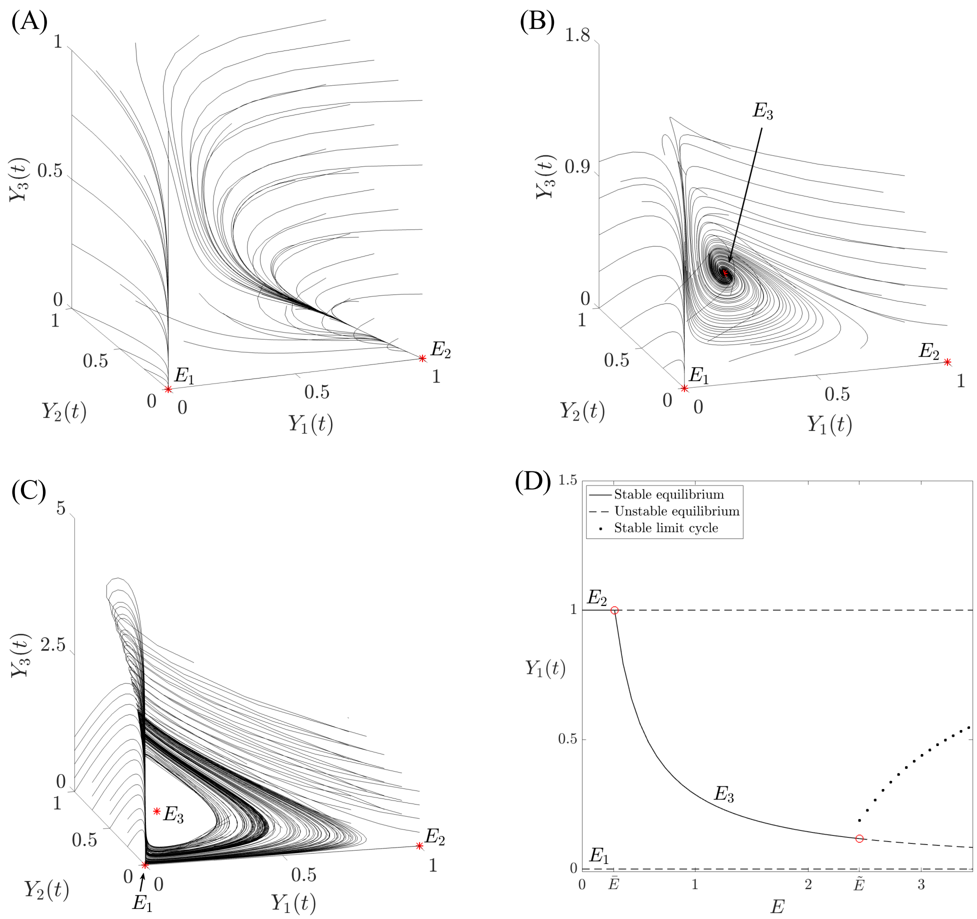


Supplementary Figure S.2: **Analysis of the dynamical system** (S.5). Phase space diagrams of the non-dimensionalized **OV module** given by equations (S.5) in three different scenarios: (A) $E < \bar{E}$, with two equilibrium points; (B) $\bar{E} < E < \tilde{E}$, with three equilibrium points; (C) $E > \tilde{E}$, where the limit cycle emerges. In (D) we show the bifurcation diagram for $Y_1$ with respect to the parameter $E$. Parameter values are set to $D = 0.58$, $F = 0.48$, and $E = 0.15$ (A), $E = 1$ (B), or $E = 3.5$ (C).

from the parameter values obtained through data fitting in Section 3.2, we observe that for all three initial virus concentration, the parameters satisfy $E > \tilde{E}$. This implies that, given enough time, the model will asymptotically approach the limit cycle around $E_3$.

In line with the classical compartmental models describing the spread of an infection, we can derive the expression of the basic reproduction number $R_0$ by applying the next generation matrix [4, 5]. Precisely, defined the matrices

$$M = \begin{pmatrix} 0 & 1 \\ E & 0 \end{pmatrix}, \qquad N = \begin{pmatrix} D & 0 \\ 0 & F \end{pmatrix}, \tag{S.8}$$

the next-generation matrix is given by $MN^{-1}$, where the spectral radius represents the basic reproduction number of the model $R_0$. Specifically, this is expressed as:

$$R_0 = \sqrt{\frac{E}{DF}}\,.$$

For the infection to spread in the tumor population, we require $R_0 > 1$, which implies that $E > DF (= \bar{E})$. In this case, the virus will be able to persist and spread. However, if $E < DF$ the only possible scenario would lead to virus extinction, as the reproduction number would not be sufficient to sustain the infection.
