## Supplementary Information S2 for "CAR T-cell and oncolytic virus dynamics and determinants of combination therapy success for glioblastoma"

### S.2 Parameter value estimations

We present the parameter values obtained from fitting the CAR T-cell module (S.1) (Table S.1), the OV module (S.4) (Table S.2), and the complete model described by system (1) in the main text (Table S.3) to the corresponding experimental data.

| E:T | $\alpha$ $(h^{-1})$ | $\theta_{TC}$ $(CI \cdot h)^{-1}$ | $\mu$ $(CI \cdot h^{-1})$ | $\delta$ $(h^{-1})$ |
|---|---|---|---|---|
| 1:10 | 0.2236 | 0.2085 | 0.026 | 0.014 |
| 1:25 | 0.2088 | 0.3965 | 0.0347 | 0.0761 |
| 1:50 | 0.1891 | 0.3640 | 0.0474 | 0.1094 |

Supplementary Table S.1: **Parameter fitting for model (S.1).** Average values of the parameters $\alpha$, $\theta_{TC}$, $\mu$, and $\delta$ from model (S.1), obtained from the fitting to the CAR T-cell and glioma cell data.

| $V_0$ [MOI] | $\alpha$ $(h^{-1})$ | $\gamma$ $(h^{-1})$ |
|---|---|---|
| 0.03 | 0.0571 | 0.0802 |
| 0.002 | 0.0508 | 0.0703 |
| 0.0008 | 0.0521 | 0.0335 |

Supplementary Table S.2: **Parameter fitting for model (S.4).** Average values of the parameter $\alpha$ and $\gamma$ from model (S.4), obtained from the fitting to the OV and glioma cell data for burst size values $b \in [0.025, 2500]$ MOI/CI. The remaining parameter are fixed to $K = 5.2246$ CI and $\omega = 0.05$ $h^{-1}$. Estimations of $\beta_{TV}$ are reported in Table S.4 and in Figure 5 in the main text.

| E:T – $V_0$ [MOI] | $\alpha$ $(h^{-1})$ | $\beta_{TV}$ $(CI \cdot h)^{-1}$ | $\theta_{TC}$ $(CI \cdot h)^{-1}$ | $\theta_{IC}$ $(CI \cdot h)^{-1}$ | $\beta_{CV}$ $(CI \cdot h)^{-1}$ |
|---|---|---|---|---|---|
| 1:50 – 0.0008 | 0.1685 | 0.0886 | 0.3263 | 0.1741 | 0.0771 |
| 1:25 – 0.0008 | 0.2458 | 0.1099 | 0.4307 | 1.0992 | 0.0924 |
| 1:50 – 0.002 | 0.2444 | 0.0529 | 0.5897 | 1.3465 | 0.0672 |
| 1:25 – 0.002 | 0.3193 | 0.0917 | 0.6293 | 3.0165 | 0.0691 |

Supplementary Table S.3: **Parameter fitting for model (1).** Average values of the parameter $\alpha$, $\beta_{TV}$, $\theta_{TC}$, $\theta_{IC}$, and $\beta_{CV}$ from model (1), obtained by fitting to the CAR T-cells, OV, and glioma cell data. The remaining parameter are fixed to $K = 5.2246$ CI, $b = 25$ MOI/CI, $\omega = 0.05$ $h^{-1}$. $\gamma$ is taken from the corresponding estimation in Table S.2, while $\delta$ and $\mu$ are taken from the corresponding estimations in Table S.1.
