## Supplementary Information S3 for "CAR T-cell and oncolytic virus dynamics and determinants of combination therapy success for glioblastoma"

### S.3 Extended analysis of burst size

Here, we complement the analysis presented in Section 3.2.1 by incorporating additional data on how variations in the burst size value $b$ influence model (S.4) parameters. Specifically, we include Table S.4, which reports the values of the parameters $\alpha$, $\beta$, and $\gamma$ for different values of $b \in [0.025, 2500]$ MOI/CI. Additionally, in Figure

| MOI \ $b$ | | 0.025 | 0.25 | 0.5 | 1 | 25 | 50 | 100 | 250 | 500 | 1000 | 2500 |
|---|---|---|---|---|---|---|---|---|---|---|---|---|
| 0.03 | $\alpha$ | 0.0414 | 0.0418 | 0.0422 | 0.0427 | 0.0456 | 0.0436 | 0.0444 | 0.0580 | 0.0639 | 0.0719 | 0.1320 |
| | $\beta$ | 3.2209 | 1.1844 | 0.8050 | 0.5312 | 0.0620 | 0.0351 | 0.0207 | 0.0101 | 0.0058 | 0.0033 | 0.0016 |
| | $\gamma$ | 0.1337 | 0.0837 | 0.0804 | 0.0782 | 0.0736 | 0.0730 | 0.0726 | 0.0722 | 0.07219 | 0.0717 | 0.0714 |
| 0.002 | $\alpha$ | 0.0653 | 0.0544 | 0.0527 | 0.0514 | 0.0486 | 0.0483 | 0.0481 | 0.0478 | 0.0477 | 0.0475 | 0.0474 |
| | $\beta$ | 3.6892 | 0.8926 | 0.5481 | 0.3303 | 0.0292 | 0.0154 | 0.0087 | 0.004 | 0.0023 | 0.0012 | 0.0006 |
| | $\gamma$ | 0.1198 | 0.0768 | 0.0731 | 0.0706 | 0.0640 | 0.0631 | 0.0624 | 0.0616 | 0.0611 | 0.0607 | 0.0602 |
| 0.0008 | $\alpha$ | 0.0580 | 0.0540 | 0.0532 | 0.0526 | 0.0511 | 0.0510 | 0.0508 | 0.0507 | 0.0507 | 0.0506 | 0.0505 |
| | $\beta$ | 4.0499 | 1.0984 | 0.6825 | 0.4155 | 0.0381 | 0.0198 | 0.0112 | 0.0052 | 0.0029 | 0.0016 | 0.0007 |
| | $\gamma$ | 0.0606 | 0.0371 | 0.0351 | 0.0336 | 0.0301 | 0.0296 | 0.0292 | 0.0288 | 0.0285 | 0.0283 | 0.0280 |

Supplementary Table S.4: **Effects of burst size variation on model (S.4) parameters.** Values of the parameters $\alpha$, $\beta$, and $\gamma$ of model (S.4) for different values of the burst size $b \in [0.025, 2500]$ MOI/CI.

S.3, we visually represent the impact of $b$ on $\alpha$ and $\gamma$, further complementing the results shown in Figure 5 of the main text. Our analysis reveals that, except for very small values of $b$ (i.e., $b = 0.025$), there is no significant effect of the virus burst size variation on the proliferative capability of tumor cells or their apoptosis rate. Finally, in Figure S.4 we illustrate the results of the analysis described in Section 3.2.2 concerning the

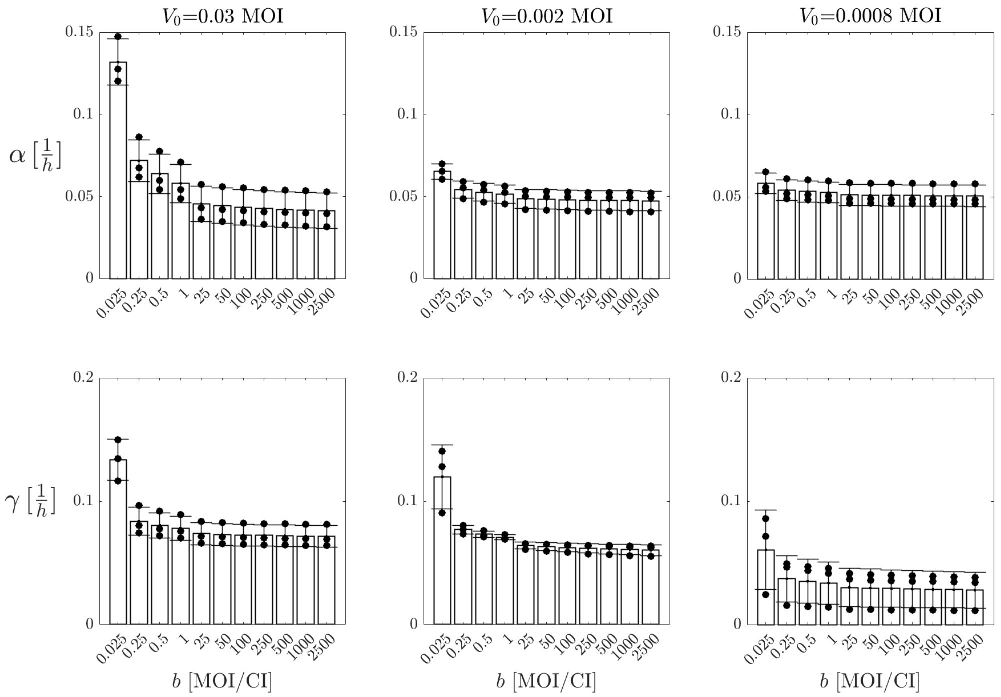


Supplementary Figure S.3: **Burst size effects on the tumor growth and apoptosis rates.** Estimation of the parameters $\alpha$ (first row) and $\gamma$ (second row) from the **OV module** given by system (S.4) for various burst size values $b \in [0.025, 2500]$. Columns correspond to the three different initial virus doses: $V_0 = 0.03$, $V_0 = 0.002$, and $V_0 = 0.0008$.

impact of the virus-specific parameters $b$ and $\omega$ on the estimation of the fractions of infected tumor cells at 24 hours (top row) and 48 hours (bottom row) obtained by using model (S.4).

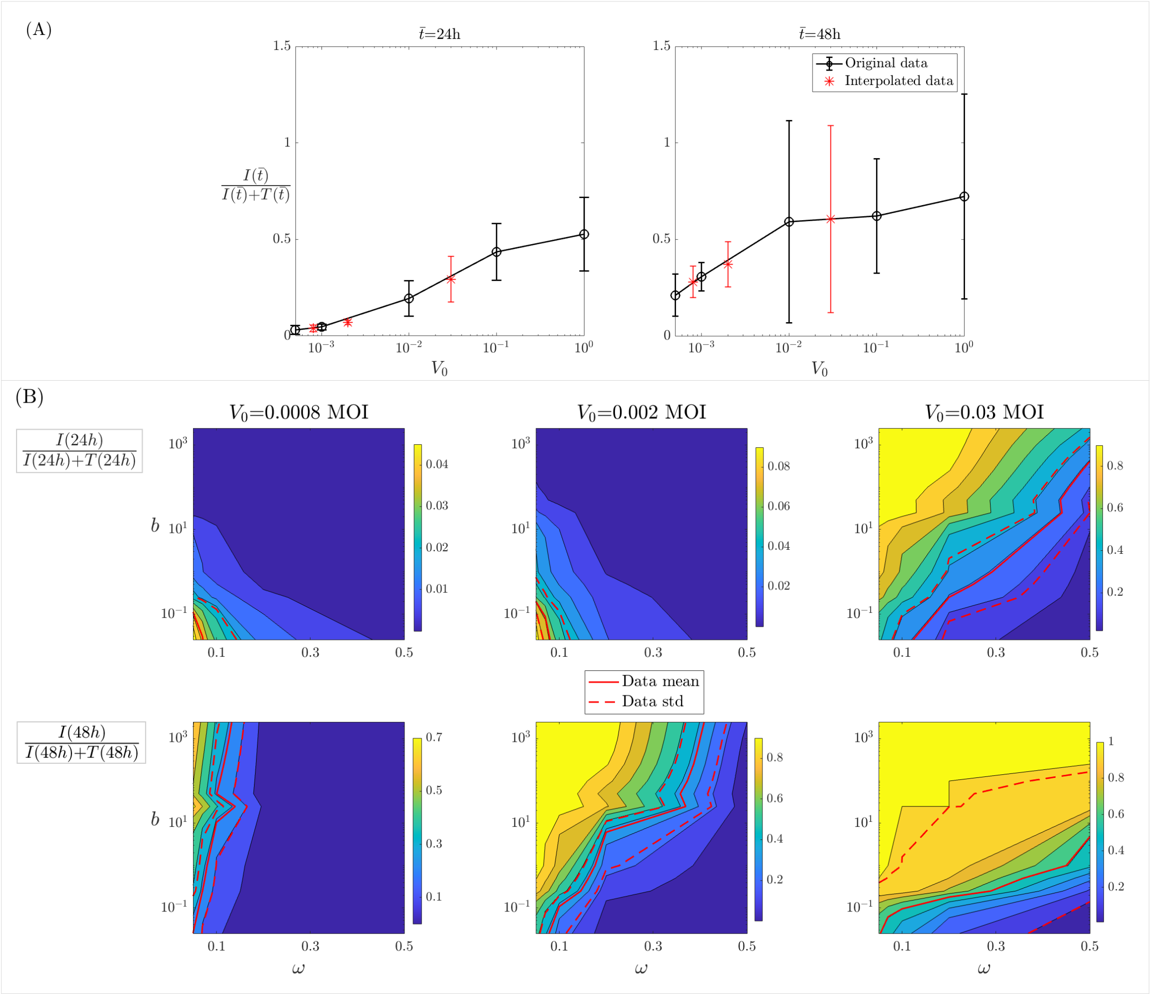


Supplementary Figure S.4: **Infected cell fraction analysis.** (A): Black circles represent experimental data for the fraction of infected tumor cells ($I(t)$) relative to the total tumor population ($I(t) + T(t)$) at 24 hours (left) and 48 hours (right) for virus doses $V_0 \in \{0.0005, 0.001, 0.01, 0.1, 1\}$. Red stars indicate the fractions of infected tumor cells at 24 hours (left) and 48 hours (right) for virus doses $V_0 \in \{0.0008, 0.002, 0.03\}$, derived through interpolation. Initial virus concentrations are presented on a log scale. Data are shown with means (markers) and standard deviations (error bars). (B): Contour plots depicting the fraction of infected tumor cells ($I(t)$) relative to the total tumor population ($I(t) + T(t)$) at 24 hours (top row) and 48 hours (bottom row). The three columns correspond to initial virus concentrations $V_0 = 0.0008, 0.002, 0.03$, respectively. The fraction of infected cells is illustrated as a function of the virus clearance rate $\omega$ (x–axis) and the burst size $b$ (y–axis, in log scale). In each subplot, the mean value of the interpolated experimental data (red markers in panel A) is represented by a red solid line, while the standard deviation (red error bars in panel A) is shown with a red dashed line.
