## Supplementary Information S4 for "CAR T-cell and oncolytic virus dynamics and determinants of combination therapy success for glioblastoma"

### S.4 Extended predictions of CAR T–OV model

Here, we complement the analysis presented in Sections 3.3.2 and 3.3.3 by incorporating additional data to evaluate the predictive capability of the complete CAR T-OV model given by system (1). Specifically, in Figure S.5, we display the qualitative behavior of the model under various initial conditions for CAR T-cells and oncolytic virus. The conditions are as follows: E:T=1:25, $V_0 = 0.0006$ MOI (A); E:T=1:50, $V_0 = 0.0006$ MOI (B); E:T=1:25, $V_0 = 0.00012$ MOI (C); E:T=1:50, $V_0 = 0.00012$ MOI (B). As anticipated, the model does not always perfectly fit the experimental data, given that we have not performed parameter estimation through data fitting. However, in some cases (panels A and B), the model exhibits a good qualitative agreement with the data. Notably, the model successfully predicts the overall tumor population dynamics over time, as shown in Figure 10 of the main text.

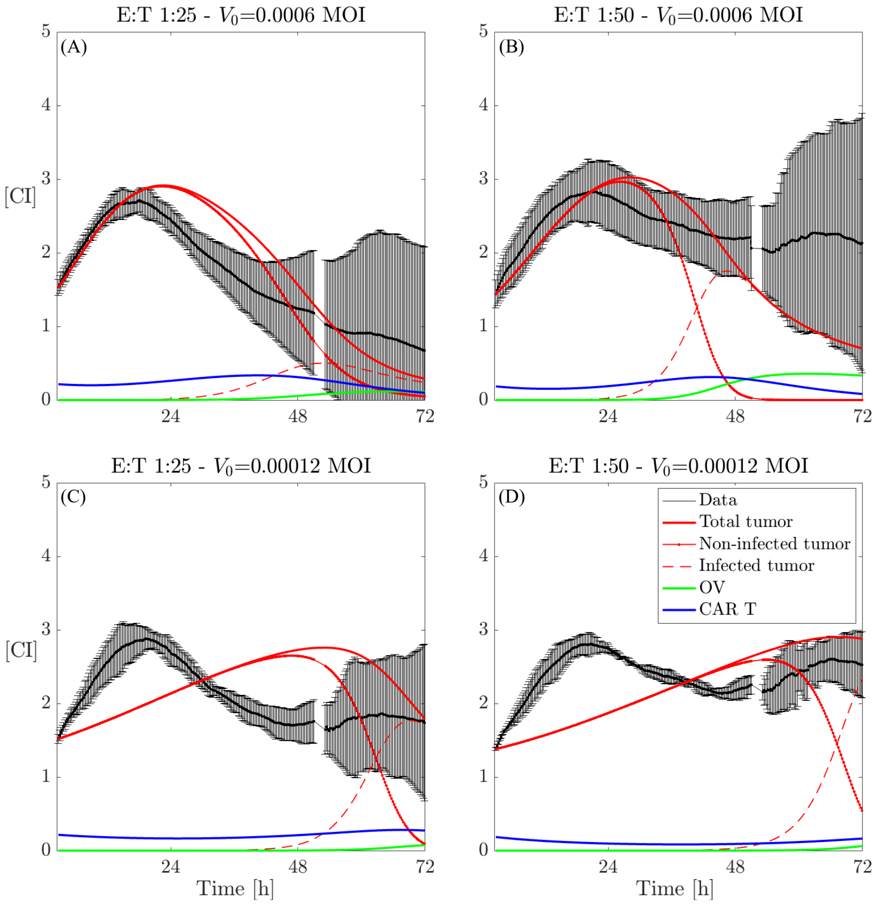


Supplementary Figure S.5: **Dynamics of model (1) and** ***in vitro*** **CAR T-cell, OV, and glioma cell data.** Dynamics of cancer cells, expressed in cell index from xCELLigence, along with the model trajectories obtained from system (1) (black: data; red continuous line: total tumor cells; red dashed–dot line: non-infected tumor cells; red dashed line: infected tumor cells; green line: OV; blue line: CAR T-cells) with initial E:T ratio and virus dose of 1:25–0.0006 MOI (A), 1:50–0.0006 MOI (B), 1:25–0.00012 MOI (C), and 1:50–0.00012 MOI (D). The parameters used in each of the four combinations of therapy are described in Section 3.3.2 in the main text.

In Figure S.6, we expand on the analysis of the effect of therapy administration delay by illustrating the potential outcomes when varying administration times are used in the two monotherapy scenarios: CAR T cell only (A) and OV only (B). The left columns in both panels display AUC values for the three different E:T ratios (top row) and $V_0$ (bottom row) for therapy delays ranging from 0 to 40 hours. The right columns present the tumor population value $T(\bar{t}) + I(\bar{t})$ at the time point $\bar{t} = 72h$, which corresponds to the end of the experiment, for the three different E:T ratios (top row) and $V_0$ (bottom row) for therapy delays ranging from 0 to 40 hours. We observe that for both monotherapy cases, no significant improvements are seen in the AUC for any of the initial E:T ratios or initial virus concentrations. Some minor improvements are noticeable only in the CAR T-cell therapy case, where a reduction in tumor population is seen at the final time point for E:T=1:50 and E:T=1:25.

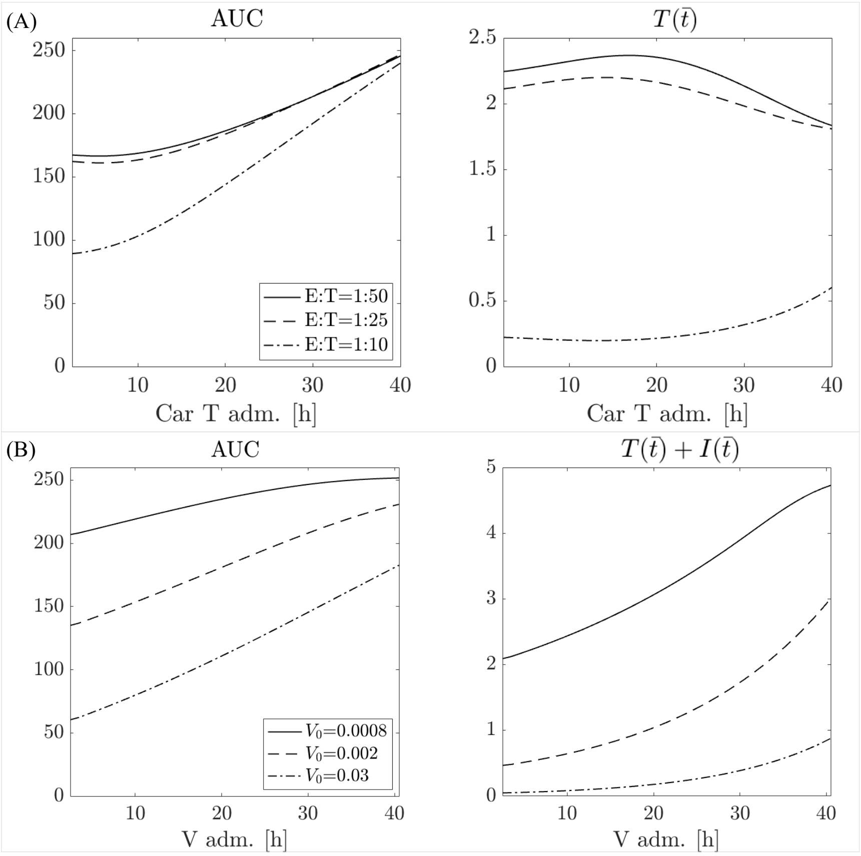


Supplementary Figure S.6: **Effects of delay in monotherapy administration.** (A): CAR T-cell monotherapy case. AUC (left column) and tumor population at the final time point $T(\bar{t})$, $\bar{t} = 72h$, (right column) calculated from model (S.1) for different therapy administration schedule. The $x$–axis represents the time of CAR T cell administration. In each plot the effect of therapy delay is shown for E:T=1:50 (continuous line), E:T=1:25 (dashed line), and E:T=1:10 (dot-dashed line). (B): OV monotherapy case. AUC (left column) and tumor population at the final time point $T(\bar{t}) + I(\bar{t})$, $\bar{t} = 72h$, (right column) calculated from model (S.4) for different therapy administration schedule. The $x$–axis represents the time of OV administration. In each plot the effect of therapy delay is shown for $V_0 = 0.0008$ MOI (continuous line),$V_0 = 0.002$ MOI (dashed line), and $V_0 = 0.03$ MOI (dot-dashed line).
